# Using light to establish habits in laboratory mice

**DOI:** 10.64898/2026.03.28.714966

**Authors:** Shu Kit Eric Tam, Xiao Xiao, Xiaoduo Cheng, Sze Chai Kwok, Benjamin Becker

**Affiliations:** Duke Kunshan University—The First People’s Hospital of Kunshan Joint Brain Sciences Laboratory, Kunshan, Jiangsu, People’s Republic of China; Division of Natural and Applied Sciences, Duke Kunshan University, Kunshan, Jiangsu, People’s Republic of China; Behavioral and Cognitive Neuroscience Center, Institute of Science and Technology for Brain-Inspired Intelligence, Fudan University, Shanghai, People’s Republic of China; The State Key Laboratory of Medical Neurobiology and MOE Frontiers Center for Brain Science, Fudan University, Shanghai, People’s Republic of China; Department of Psychology, The University of Hong Kong, Hong Kong, People’s Republic of China; SRT AI, Society and Social Dynamics, The University of Hong Kong, Hong Kong, People’s Republic of China

**Keywords:** Animal models, Mice, Instrumental conditioning, Habit formation, Light, Digital devices

## Abstract

**Background and aims:** Perseverative behaviours are commonly assessed using operant paradigms in which rodents work for drugs or food under physiological deprivation, limiting translational relevance to some behavioural addictions. Here we validated an operant paradigm in which the acquired behaviour is driven neither by physiological needs nor hedonic responses.

**Methods:** Mice were trained to lever-press for green light. Exp.1 used a within-subjects design to examine lever discrimination and whether responding could be **“**satiated**”** by light preexposure. Exp.2 examined instrumental contingency using a between-subjects design, with light delivery equated between contingent and non-contingent groups. Exp.3 replaced green light with dim red light producing less retinal photoreceptor excitation but comparable heat to assess non-photic cues. Exp.4 examined whether green light could affect food seeking different motivational states.

**Results:** In Exp.1, green light supported lever discrimination. Among high responders, the satiation effect was modest (<15% reduction) and did not deter lever pressing. In Exp.2, instrumental contingency promoted response acquisition whereas random light delivery did not. In Exp.3, dim red light failed to sustain behaviour, producing ∼50% response decrement. In Exp.4, light potentiated food seeking under *ad libitum* feeding.

**Discussion and conclusions:** Response-contingent light serves as a reward to establish operant responding, which cannot be explained by alerting effects or thermal cues. Our study bridges the gap between animal models and findings from humans that coloured light may exacerbate smartphone use and that light therapy may reshape reward circuits in individuals with Internet gaming disorder symptoms [Li et al. (2026) *Advanced Science* **13:**e14044].

## Introduction

In the broadest sense, everyday habits are perseverative behaviours that can be observed repeatedly under certain situations, and they take time to modify (Hull, 1943; Bouton, 2024). Experimental studies investigating the mechanisms of habit formation and compulsive behaviours have been conducted in humans and laboratory animals for decades. While habits are mostly adaptive and essential for survival, they can be formed in response towards a multitude of primary and conditioned reinforcers (Robbins et al., 2024). Traditionally, everyday habits are characterised in terms of goal directedness—whether the behaviour under investigation is sensitive to reinforcer values driven by *response* → *reinforcer* associations, or simply an automatically elicited outcome-insensitive response driven by *stimulus* → *response* associations (Robbins et al., 2024). This dichotomy, however, is likely to be oversimplistic, as both mechanisms contribute to clinically maladaptive behaviour (Hogarth, 2020; Robbins et al., 2024). Regardless of theoretical standpoints, pathological habit formation and excessive execution are transdiagnostic features of a group of related clinical conditions characterised by compulsions, including obsessive–compulsive disorder, substance use disorders, and behavioural addictions (Klugah-Brown et al., 2021; Robbins et al., 2024). Although these conditions share behavioural and neural features in humans, the corresponding animal models are often disorder specific rather than developed with a transdiagnostic focus. For example, animal models for substance use disorders may not be directly relevant to behavioural addictions such as gambling disorder (and *vice versa*), as the specific reinforcers driving these behavioural phenotypes are fundamentally different (Spanagel, 2017; Zentall, 2023).

In traditional operant paradigms for both rats and mice, habits are induced using instrumental (operant) conditioning paradigms, in which an animal learns to self-administer a reinforcer that satisfies a physiological need or elicits a hedonic response. Typical reinforcers include caloric food or water delivery (Reinagel, 2018; Chevée et al., 2023), intravenous substance infusion (Leonardo et al., 2023; Marini et al., 2025), and intracranial reward-circuit stimulation (Wise & McDevitt, 2018; Wolff & Saunders, 2024). These paradigms provide experimental models to investigate processes underlying compulsions and addictive behaviours triggered by food and drugs as well as their associated cues. However, these operant paradigms do not fully capture processes underlying maladaptive behaviours triggered by *non-food*, *non-drug* primary reinforcers, particularly those relevant to obsessive–compulsive behaviours and behavioural addictions including digital technology-based disorders (Klugah-Brown et al., 2021; Robbins et al., 2024; Tam et al., 2025).

To circumvent this limitation, the mouse *operant sensation seeking* paradigm can be utilised (Olsen & Winder, 2009; Gancarz et al., 2023). Unlike traditional paradigms, the reinforcer in this paradigm is not food or drug but a multimodal sensory stimulation, comprising a compound cue with sound and light of varying duration, flickering frequency, and spatial position to maintain novelty of the sensory reinforcer (Olsen & Winder, 2009; Gancarz et al., 2023), thereby maintaining the animal’s engagement with the operant task. This paradigm is effective in establishing operant responding, requiring at least 6 hours to extinguish completely within a single extinction session (Olsen & Winder, 2009). Although effective, it is unclear which component of the sensory complex reinforces the operant response—it could be light, novelty, complexity, or uncertainty of the sensory reinforcement (or *all* these factors). Thus, in addition to determining an animal model for habit and compulsive behaviour, our research was motivated by the continuing interest in the diverse roles of artificial light in brain health and disease (Huang et al., 2024; Mahoney et al., 2024). We therefore systematically investigate the reinforcing property of light, isolating this from novelty and complexity of the sensory reinforcement in the operant sensation seeking paradigm (Olsen & Winder, 2009; Gancarz et al., 2023). Specifically, we show that unimodal, monochromatic light—a cue that is *neither* physiologically significant *nor* physically salient—is sufficient to establish a perseverative operant response in laboratory mice.

Using discrete light cues as reinforcers has a major advantage over other non-food, non-drug reinforcers, such as thermal cues (Gordon et al., 1998), novel objects (Bevins & Basher, 2005), running wheels (Nishitani et al., 2025), and social interaction with conspecifics (Ramsey et al., 2023). Notably, the dose of light administration (e.g., light duration and wavelength) as well as timing and instrumental contingency (i.e., the causal relationship between the preceding action and light delivery) can be precisely controlled by the experimenter. This allows precise adoption of traditional schedules of reinforcement (ratio or interval schedule) in our study. Wildtype mice (C57BL/6J) instead of rats are chosen as the model organism, because rats exhibit decline in light self-administration within a session and lack noticeable improvement across days (Glow & Russell, 1973; Lloyd et al., 2014), due to rapid reinforcer habituation under *fixed*-ratio (FR) schedules (Lloyd et al., 2014). This is not the case for mice in our current study. Here we use male mice to demonstrate the reinforcing effectiveness of light, because in our pilot experiment male mice responded more on levers than female mice from the same litter (therefore of the same age) due to differences in body weight and physical strength (Tam, 2024).

We use two distinct environments to examine if light self-administration can be affected by training context, given the influence of context on drug self-administration in rodents. Specifically, the novelty of the test environment in which rats are assessed can facilitate acquisition of cocaine self-administration but not saline self-administration (Caprioli et al., 2007); this indicates the potential contribution of contextual factors to maladaptive habit formation. Accordingly, we include two training environments (the standard operant chamber and Y–maze) differing in their size, geometry, and material: the size and geometry of the operant chamber resemble the mouse’s home cage, whereas the Y–maze is longer and has a distinct and novel geometrical layout (**Figure 1*a***).

**Figure 1.**
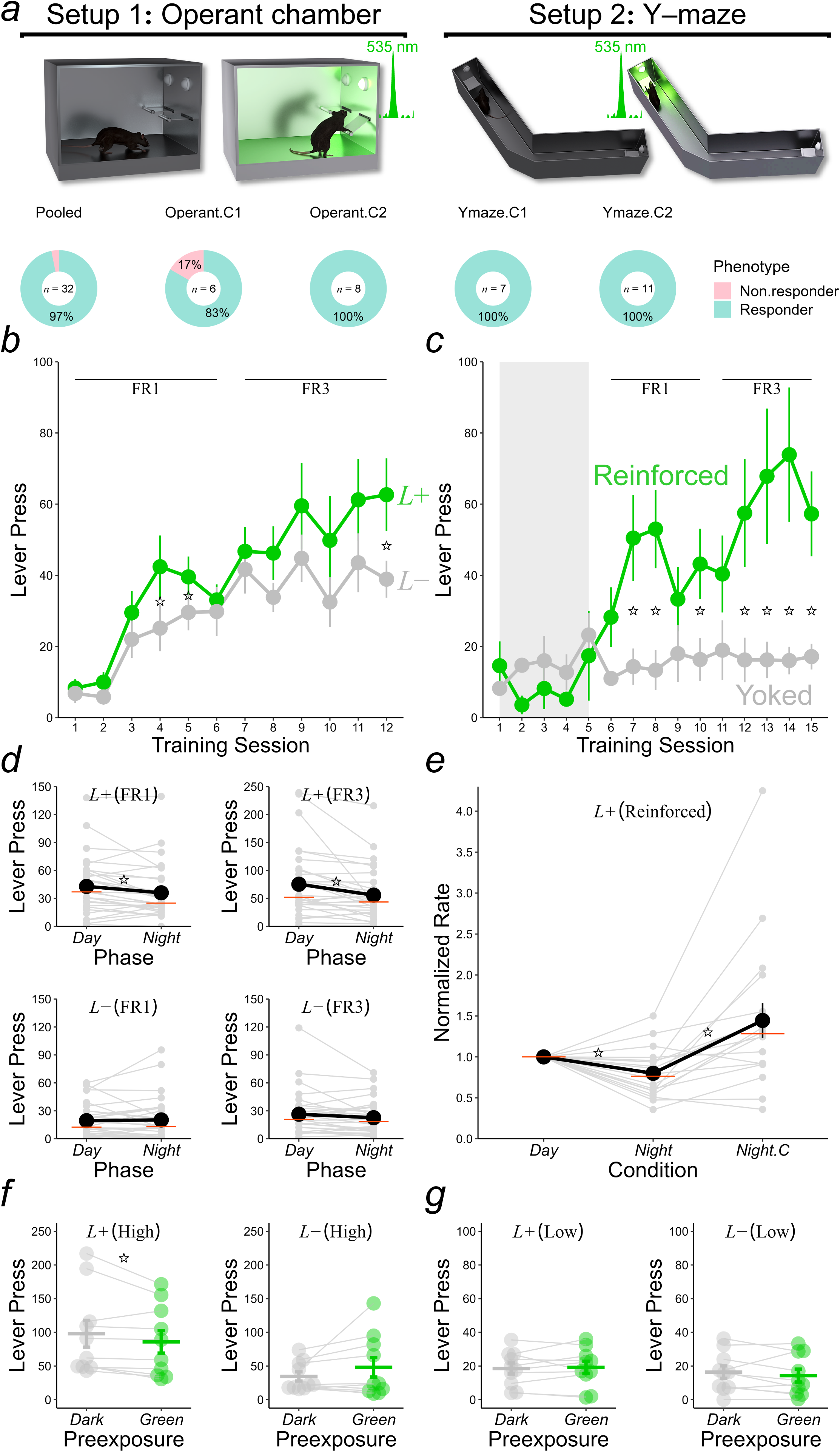
| Operant light self-administration under FR schedules. (***a***) Two training environments were used. Pie charts show proportions of Responders and Non-responders in all cohorts of mice that received operant training (Experiment 1: *Operant*.*C1*, *Operant*.*C2*, and *Ymaze*.*C1*; Reinforced mice in Experiment 2: *Ymaze*.*C2*). One mouse in *Operant*.*C1* stopped responding in the last two FR3 day sessions, and it was classified as a non-responder. (***b***) In Experiment 1, green light supported two-lever discrimination (pooled across cohorts *Operant*.*C1* and *Ymaze*.*C1*). *L*+ (*green*) represents the lever reinforced according to fixed-ratio schedules (FR1 and FR3); *L*− (*grey*) represents the inactive lever without consequence. (***c***) In Experiment 2, instrumental contingency between lever pressing and light delivery was required for light self-administration acquisition (Reinforced mice; cohort *Ymaze*.*C2*), as there was no change in lever responding in control mice receiving random light presentations (Yoked Control mice). Sessions 1–5 shaded in *grey* were conducted in darkness without reinforcement schedules. (***d***) Under FR1 and FR3, mice showed reduced *L*+ responding at night (pooled across all cohorts); the time-of-day effect was lever specific (Time of day × Lever interaction: *p* = 0.0032). (***e***) The decline in responding at night was reinstated by lengthening light duration from 2.5 s to 4 s in *Operant*.*C2* and from 0.5 s to 2.5 s in *Ymaze*.*C2* (condition *Night*.*C*). Means, medians, and individual data are coloured in *black*, *red*, and *grey*, respectively, in panels ***d*** and ***e***. (***f***, ***g***) Mice in the High-responding subgroup showed a modest satiation effect on *L*+ after 30-min green light preexposure; this effect was not detected in the Low-responding subgroup presumably due to a floor effect (Preexposure Condition × Lever × Responder Subgroup interaction: *p* = 0.019). Significant differences (*p* < 0.05) are denoted by star symbols (*).

## Methods

### Animals and housing condition

A total of 52 male C57BL/6J mice were purchased from a local supplier (Model Organisms, Shanghai, China). In Experiments 1–3, initial sample sizes in light-training cohorts *Operant*.*C1*, *Operant*.*C2*, *Ymaze*.*C1*, *Ymaze*.*C2*, and the control group were as follows: *n* = 6, *n* = 8, *n* = 8 (one died due to fighting during transportation prior to arrival at the colony), *n* = 11, and *n* = 11 (one was found dead before the experiment commenced); whereas the size of the food-training cohort in Experiment 4 was *n* = 8. Cohort sizes in light-training groups were comparable to our pilot study that demonstrated increment of light-seeking response with *n* = 8 (Tam, 2024), which was also comparable to typical cohort sizes in published operant food-seeking experiments (e.g., 4–8 mice per group in Shaw et al., 2004) and operant intracranial self-stimulation experiments (e.g., 4–6 mice per group in Rossi et al., 2013). All mice in this study were 8 weeks old upon arrival at the animal facility. In all experiments, they were housed individually (for home cage activity monitoring) under a 12-h light:12-h dark cycle [Zeitgeber time (ZT)00 started at 06:00 and ZT12 started at 18:00] with *ad libitum* access to food and water; in Experiment 4, 8 mice were maintained at 85–90% of their free-feeding body weights for 8 days while learning the food self-administration task. Corn cob bedding was used as bedding material in the mouse cage; four cotton balls were placed inside the cage for the mouse to use as nesting material. Water bottles were changed once every 10 days, and cage bedding was changed once every 3 weeks. Mouse cages were kept inside enclosed wooden chambers; for 6 out of 12 mice in Experiment 1 and 6 out of 8 mice in Experiment 4, passive-infrared sensors (PIR) were placed 35 cm above their cages to monitor activity in 10-s time bins (Tam et al., 2021). Cool white LED lights in the chamber provided animals with 100–200 lux light in the light phase. The spectral power distribution of the white LED comprised a higher, narrower peak at 455 nm with a lower, broader peak at 570 nm, comparable to that in our previous study (Tam et al., 2021). The ambient temperature in the animal holding room was maintained at 21 ± 1.5°C. Operant training commenced one week after animals’ arrival at the colony. During the acquisition phase of light and food self-administration, operant sessions were conducted in the light phase at ZT01–04 *and* in the dark phase at ZT13–16. Training times are summarised in **Supplementary Table 1*a***. Experimental sessions were conducted seven days a week by one experimenter (S.K.E.T.; except that the last probe session in Experiment 3 was conducted by X.C.). Behavioural setups were in the same room where animals were housed, so that the delay between picking up the mouse from its home cage and placing it into the behavioural setup was minimal (usually <1 min)

### Setup 1: Operant chamber

The operant chamber (16 cm × 14 cm × 13 cm; ENV-307A, Med Associates, Vermont) was placed inside an enclosed chamber (59.5 cm × 35.5 cm × 37.5 cm) with a ventilation fan, which remained off during the session. The operant chamber comprised two short stainless-steel walls and two long transparent plastic walls (the front one was the door); the ceiling was made of transparent plastic. The floor grid comprised 19 stainless steel bars spaced 0.5 cm apart; each bar had a diameter of 0.3 cm, running parallel to the short walls. A tray with tissue sheets was placed below the floor grid to collect urine and droppings from the mouse; this was cleaned after every session. A receptacle with an opening of 2.7 cm × 2.1 cm and a depth of 2 cm was located on one of the short walls, equidistant from the long walls and 0.2 cm above the floor grid. On each side of the receptacle, there was one stainless steel lever (with a paddle surface of 1.6 cm × 0.8 cm and 2.3 cm above the floor grid) that was inserted into the chamber at the start and retracted at the end of the light self-administration session. The force required to activate the response lever was ≥2.5 g. Lever pressing responses were recorded and reinforcement schedules were delivered by Med-PC (version V, Med Associates, Vermont).

### Stimuli in the operant chamber

In the first light self-administration experiment (**Figure 1*b***), the light reinforcer was 4-s continuous illumination of the monochromatic green LED house light, located 10.5 cm above the floor grid on the short wall opposite to the receptacle. A spectrophotometer (XL-500 BLE, nanoLambda, Korea) was used to measure the spectral power distribution of the green LED (*λ*_max_ = 535 nm), which is shown in **Figure 3*c***. No food was delivered in the light self-administration experiment. In the food self-administration experiment (**Figure 4**), the food reinforcer was delivery of one 20-mg cream-flavoured pellet (2.5 mm in diameter) with an ingredient composition of ≥10% protein, ≥5% fat, and <10% carbohydrate, and an energy content of 4–5 kcal g^−1^ (Youer Equipment Scientific, Shanghai). The food pellet was delivered directly into the base of the receptacle by a pellet dispenser. For both experiments, the operant chamber remained in complete darkness except during presentation of the light reinforcer.

**Figure 2.**
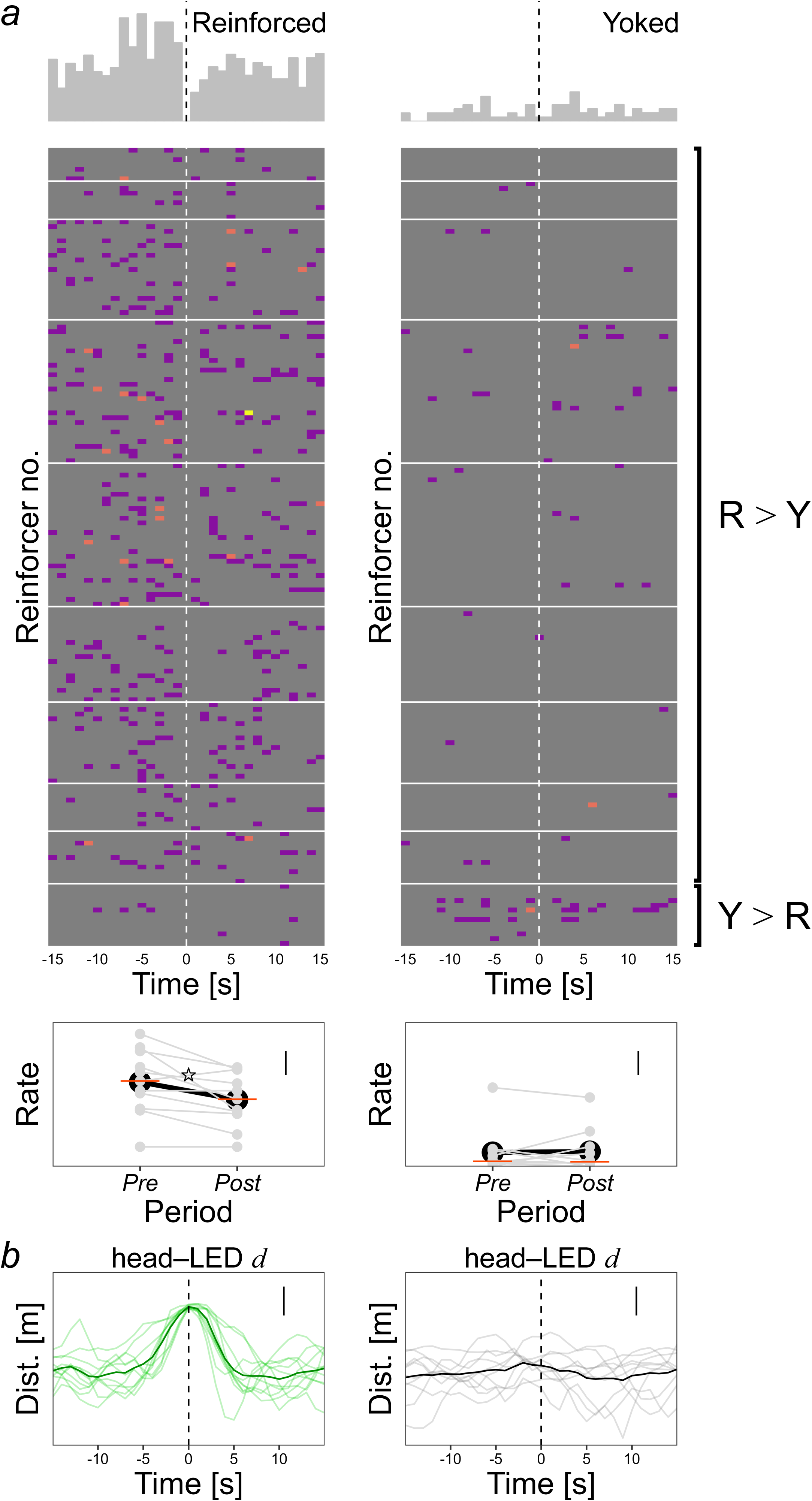
| Lever responding during the 15-s period preceding (*pre*-LED) and following light onset (*post*-LED) under FR5 and proximity to the LED location. (***a***, *top* and *middle*) Histograms and raster plots show local response rates (s^−1^) at the group level and trial by trial during *pre*-LED and *post*-LED periods in Reinforced and Yoked Control mice. Each row in the raster plot show delivery of one reinforcer. Records from individual mice were stacked and aligned to bins containing lever presses that led to reinforcer delivery, which was defined as *time* = 0 s (*vertical dashed* lines). *White horizontal lines* demarcate records from different Reinforced–Yoked Control pairs. Bins coloured in *purple*, *red*, and *yellow* represent 1, 2, and ≥3 lever presses s^−1^, respectively, whereas *grey* bins represent no lever press. Reinforced mice showed higher response rates than Yoked Control mice before and after light onset, except the last pair with the opposite pattern. (***a***, *bottom*) Average response rates (s^−1^) declined during the *post*-LED period relative to the *pre*-LED period in Reinforced mice (\**p* < 0.05), consistent with the typical post-reinforcement pause under appetitive FR schedules (Rickard et al., 2009). Means, medians, and data from individual mice are coloured in *black*, *red*, and *grey*, respectively. Vertical scale bars represent 0.025 press s^−1^. (***b***, *left*) Shortly after light onset, Reinforced mice moved away from the lever and oriented towards the LED, as indicated by a rapid drop in the distance between the LED location and mouse’s snout (head–LED *d*). Lever–light alternation was seen in all mice in the Reinforced group (see also **doi.org/10.6084/m9.figshare.31768912**). *Thin green* lines show individual data, and the *thick green* line shows the group average. (***b***, *right*) By contrast, there was no change in the head–LED distance in Yoked Control mice, indicating that they did not alternate between lever pressing and orienting towards the LED location during the *post*-LED period. *Thin grey* lines show individual data, and the *thick black* line shows the group average. Vertical scale bars represent 0.05 m (lever–LED distance ≈ 0.34 m).

**Figure 3.**
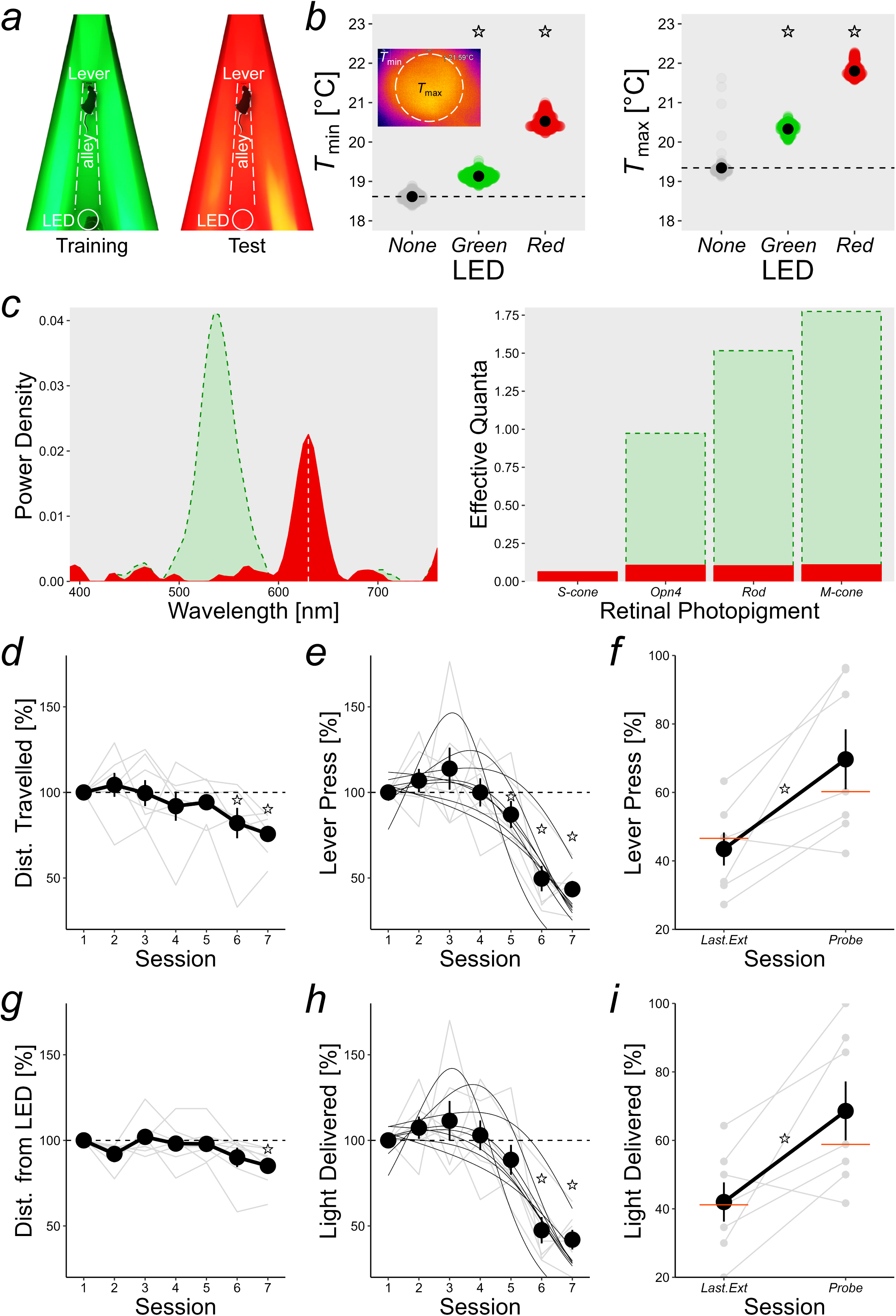
| **Response decrement under FR5 with dim red light.** (***a***) After cohort *Ymaze*.*C1* were retrained with green light, it was replaced with dim red light. (***b***) Minimal and maximal surface temperatures (*T*_min_ and *T*_max_, respectively) of the green and red LED lights as measured by thermal imaging (see Methods: *Y–maze*) are shown. The average LED surface temperature was higher when either LED light was illuminated than when it was off (*None*: 18.98 ± 0.016°C; *Green*: 19.73 ± 0.006°C; *Red*: 21.16 ± 0.006°C; \**p* < 0.005 *versus* no-light condition). (***c***, *left*) Spectral power distributions of the monochromatic green and red LED lights were measured (in W⋅m^−2^⋅nm^−1^; range: 390–760 nm in 5-nm bins) using a spectrophotometer. LED peak emission wavelengths were at 535 nm and 630 nm (indicated by the vertical dashed white line). (***c***, *right*) Effective quantal values consider each photopigment’s spectral sensitivity and describe the amount of light (in photopic lux) needed to produce a certain photopigment-specific (α-opic) illuminance value (Lucas et al., 2014, 2024; Peirson et al., 2018). α-opic irradiances of green and red LED lights for the mouse short-wavelength sensitive cone opsin (*S*-cone), melanopsin (*Opn4*), rhodopsin, and medium-wavelength sensitive cone opsin (*M*-cone) were approximated using the Rodent Toolbox (Peirson et al., 2018; Lucas et al., 2024) The dim red LED light activated the four photopigments in the mouse retina much less than the green LED light. (***d***, ***e*** and ***g***, ***h***) Sessions were conducted with red light delivered according to FR5. Behavioural measures included distance travelled (panel ***d***), lever responding (panel ***e***), head–LED distance (panel ***g***), and number of reinforcer delivery (panel ***h***). Significant differences from the first red-light session are denoted by star symbols (\**p* < 0.05). For data in panels ***e*** and ***h*** where there were orderly changes across days, a 3-paramater sigmoid function, *y* = *a***e**^−*x*^/[1+**e**^−*b*(*x*−*c*)^], was fitted onto every individual extinction curve using nonlinear least-squares estimates obtained from the `nls` function. (***f***, ***i***) The day after the last extinction session, a test was conducted not by the original experimenter (S.K.E.T.) but by a different experimenter (X.C.), providing an extra-experimental context that was different from all previous sessions. Non-specific changes in the experimental context, potentially including handling cues (Hurst & West, 2010) and experimenter-related odours (Sorge et al., 2014), caused recovery of the operant response (*Last*.*Ext versus Probe* \**p* < 0.05), resembling the context specificity of extinction in appetitive tasks (Bouton, 2024). However, recovery was partial as operant responding and number of reinforcers collected were still below 100% [69.67% ± 8.76% *versus* 100%: *t*(6) = −3.46, *p* = 0.013 (2-tailed) in panel ***f***; 68.58% ± 8.62% *versus* 100%: *t*(6) = −3.65, *p* = 0.011 (2-tailed) in panel ***i***]. In panels ***d***–***i***, means, medians, and data from individual mice are coloured in *black*, *red*, and *grey*, respectively; data were normalised to initial values in the first red-light session.

**Figure 4.**
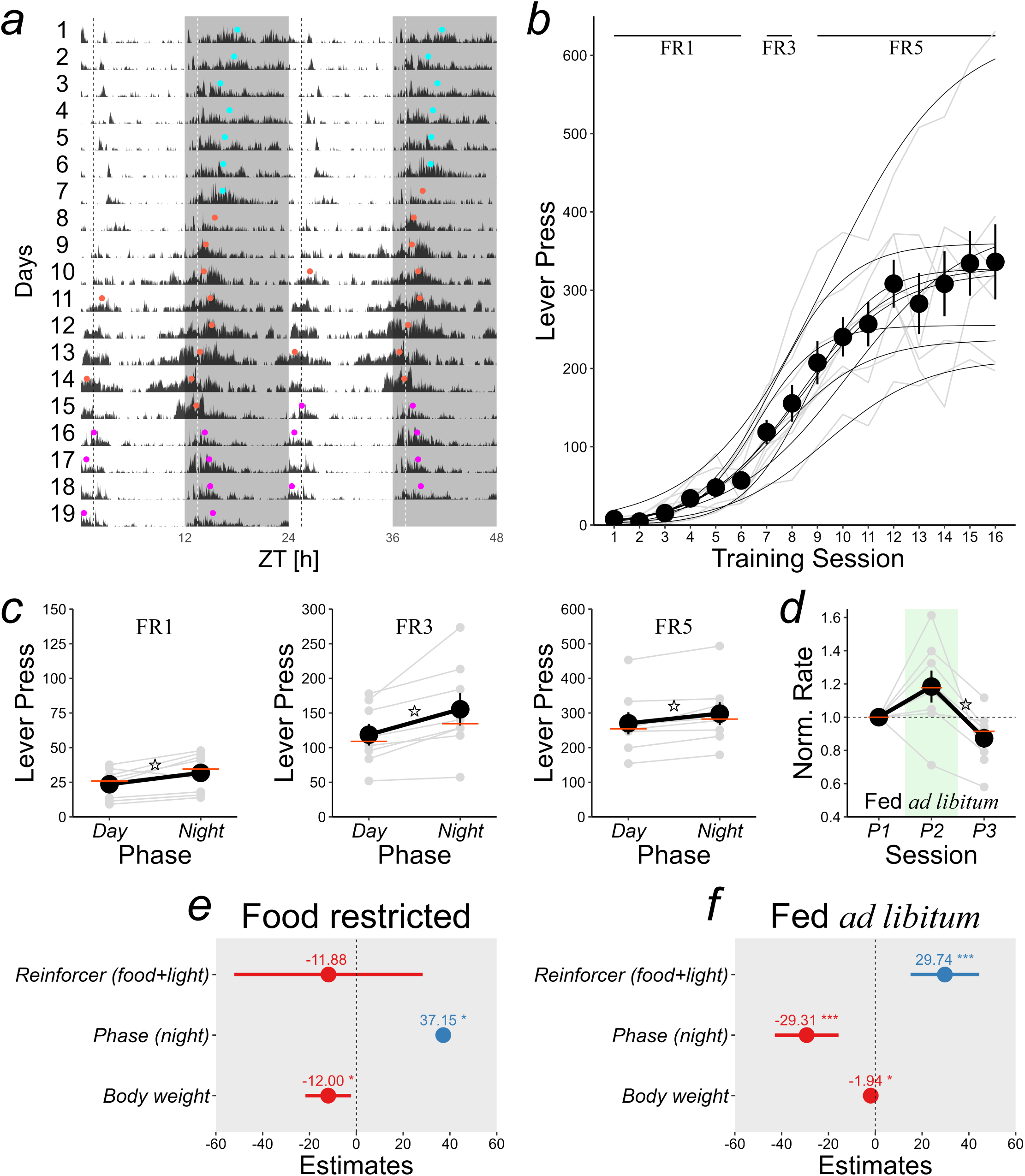
| **Light potentiates operant responding for food.** (***a***) The double-plotted actogram (data pooled across mice) shows home cage PIR activity (*black*) in Experiment 4. The dark phase of the 12-h light:12-h dark cycle is shaded in *grey*. Coloured dots represent activity midpoints, defined as the time points at which total activity in the preceding 8 h and activity in the subsequent 8 h were equivalent (see Methods: *Data analyses*). ●: midpoints before the experiment began (days 1–7); ●: midpoints during operant training under food restriction (days 8–15); ●: midpoints during operant testing with food *ad libitum* in home cages (days 16–19). Vertical dashed lines indicate operant training and testing times on days 8–19, during which diurnal rhythm was disrupted (see **Supplementary** Figures 4a,***b***). (***b***) Acquisition of food self-administration under FR1 (training sessions 1–6), FR3 (training sessions 7 and 8), and FR5 schedules (training sessions 9–16); mice were trained twice per day. A 3-paramater sigmoid function, *y* = *a*/[1+**e**^−*b*(*x*−*c*)^], was fitted onto every individual learning curve using nonlinear least-squares estimates obtained from the `nls` function. (***c***) Mice responded more at night under all ratio schedules in training sessions 1–16 (\**p* < 0.05). (***d***) When mice were fed *ad libitum*, a 2.5-s green light cue that was delivered in conjunction with food potentiated operant responding under FR5 in the second probe session (\**p* < 0.05, comparing probe session *P2* with food and light *versus* probe session *P3* with food only). (***e***, ***f***) Fixed-effect estimates (*slopes*) and 95% confidence intervals (CIs) of the relative contributions of reinforcer summation (food and light relative to food only), time of testing (at night relative to the light phase), and body weight to operant responding under food restriction (during FR5 sessions 9–16) and under food *ad libitum* (during FR5 probe sessions *P1*–*P3*). Positive effects with *slopes* > 0 and negative effects with *slopes* < 0 are represented in *blue* and *red*, respectively. Asterisks indicate significant effects in linear mixed-effects models. (e) Under food restriction, night testing had a positive effect (*χ*^2^ = 5.69, *p* = 0.017) and body weight had a negative effect (*χ*^2^ = 5.55, *p* = 0.018), whereas adding light to food delivery had no effect (*χ*^2^ = 0.34, *p* = 0.56). (f) When mice were fed *ad libitum*, body weight still affected operant responding negatively (*χ*^2^ = 5.91, *p* = 0.015) but the effect was attenuated; its slope estimate dropped from −12.00 [95% CI: −21.36, −2.17] to −1.94 [95% CI: −3.65, −0.23]. Adding light to food delivery had a positive effect on operant responding (*χ*^2^ = 13.91, *p* = 0.00019), consistent with panel ***d***. Light potentiation was not driven by any Pavlovian-conditioned incentive due to a lack of forward *light* → *food* pairings (cf. Wheeler et al., 2008) before the test.

### Setup 2: Y–maze

To examine if spatial complexity of the environment would affect operant choice behaviour guided by the light reinforcer, we also conducted operant training in the Y–maze. The maze (SansBio, Jiangsu, China) was made of white acrylic. It comprised a central triangular area (15 cm^2^) connecting to three rectangular arms (33.7 cm long and 5 cm wide) separated by 120°; walls of the maze were 20 cm high. The central area and the three arms could be partitioned into separate areas by white acrylic panels; the wall at the end of each arm was also a removable white acrylic panel. One of the three arms was the start arm. Shortly before the start of each session, the mouse was put into the end of the start arm. As soon as the animal ran into the central area, the partition behind it was lowered manually so that it could not return to the start arm during the session. Stainless steel levers (80120MM, Campden Instruments, England) were inserted into the end of the other two arms. Each lever had a paddle surface of 1.6 cm × 0.8 cm and was 2.5 cm above the floor; it was non-retractable and remained in the Y–maze during the session. The force required to activate the response lever was ≥2.5 g as in the operant chamber. A camera with infrared LED lights were positioned 100 cm above the Y–maze. Real-time tracking of exploratory activity was conducted in ANY-maze (version 7.49, Stoelting, Illinois), tracking the mouse’s head position every 100 ms. The timing of lever pressing was recorded, and reinforcement schedules were delivered by ANY-maze *via* the AMi-2 digital interface (60064, Stoelting, Illinois). After the completion of the experiment, to quantify the amount of heat generated by the green and red LED lights in the arm of the Y–maze (**Figure 3*a***), a thermal imaging camera (Xi 400, Optris, Berlin, Germany) was positioned 5 cm from the LED surface to record surface temperature while each LED was illuminated continuously for 10 min (**Figure 3*b***).

### Stimuli in the Y–maze

In the first light self-administration experiment (**Figure 1*b***), the accessible areas included the central area and two arms equipped with response levers, but not the start arm (hence it was depicted as a V–maze rather than a Y–maze in **Figure 1*a***). The light reinforcer was 4-s continuous illumination of the monochromatic green LED light located 18.5 cm above the floor in the central area. The spectral power distribution of the green LED (*λ*_max_= 535 nm) is shown in **Figure 3*c***. Just like in the operant chamber, the maze remained in complete darkness except during presentation of the light reinforcer. In the second light self-administration experiment, one mouse from the Reinforced group and one from the Yoked Control group were run *simultaneously*. Each mouse was confined to one arm with a single response lever. In each arm, a green LED light was positioned 18.5 cm above the floor on the wall opposite to the response lever; the central area and start arm were inaccessible. Whenever the Reinforced mouse received the light reinforcer (with a duration of either 0.5, 2.5, or 4 s; see **Supplementary Table 1*b***), the green LED lights in *both* arms were presented at the same time, so that light exposure was perfectly matched in every Reinforced–Yoked Control pair. In the response-decrement experiment (**Figure 3**), animals were trained and tested in the right arm only, and the green LED light was in the central area opposite to the response lever (see example video clip: doi.org/10.6084/m9.figshare.31768912); the duration of the light reinforcer was 4 s. A dim red LED light (*λ*_max_ = 630 nm) was used as the reinforcer the response-decrement experiment; its spectral power distribution and the extent to which it excited different mouse retinal photoreceptors are shown in **Figure 3*c***.

### Procedure in Exp.1: Two-choice discrimination and satiation test (n = 21)

In the first experiment (**Figures 1*a***,***b***), lever pressing on *L*+ but not *L*− was reinforced under fixed-ratio (FR) schedules of reinforcement; positions of *L*+ and *L*− were counterbalanced across mice. FR1 and FR3 were programmed on *L*+ in sessions 1–6 and sessions 7–12, respectively. Light duration was 4 s for cohorts *Operant*.*C1* and *Ymaze*.*C1*, and it was 2.5 s for cohort *Operant*.*C2*. In the operant chamber, a session was terminated when 45 min elapsed, or 30 light reinforcers were delivered; in the Y–maze, a session was terminated when 30 min elapsed, or 30 light reinforcers were delivered. After acquisition, test sessions were conducted to examine the effect of preexposure to darkness *versus* green light on subsequent operant responding. During dark and continuous green light preexposure, mice were left in the operant chamber (with lever retracted) or in the start arm of the maze (where no lever was present) for 30 min; 30-min light exposure exceeds the total light duration (≤2 min) a mouse can obtain in an operant session. At the end of the 30-min period, operant sessions began as usual with both levers inserted into the operant chamber or with the partition between the central area and start arm raised manually to let the mouse enter the test arena. Each mouse received *both* darkness and green light preexposure conditions. The order of preexposure was counterbalanced, so that some animals received darkness and green light preexposure on the first and second test sessions, respectively, whereas remaining animals received the opposite arrangement (**Supplementary Table 1*a***).

### Procedure in Exp.2: Instrumental contingency versus random light delivery (n = 21)

In the second experiment (**Figure 1*c***), sessions 1–5 were conducted in darkness without any reinforcement schedule to assess baseline responding in a subset of Reinforced and Yoked Control animals (**Supplementary Table 1*a***). FR1 and FR3 were programmed on the lever in sessions 6–10 and sessions 11–15, respectively; from session 16 onwards, animals were tested under FR5 (**Figure 1*e***). During initial training, identical reinforcement schedules were used in the two replications (**Supplementary Table 1*a***) but light duration differed (**Supplementary Table 1*b***): For Reinforced (*n* = 5) and Yoked Control animals (*n* = 4) in the first replication, light duration was 4 s; for Reinforced (*n* = 6) and Yoked Control animals (*n* = 6) in the second replication, it was 2.5 s. As in Experiment 1, light-duration manipulation was incorporated into the experimental design, as operant responding is known to be sensitive to food and drug reinforcer magnitude (Reed & Wright, 1988; Spear & Katz, 1991; Rickard et al., 2009). By varying light duration within a limited range while controlling for all other factors (including housing condition, age, and training time), this allows us to examine the generalizability of light-seeking behaviour. Despite using different light durations in initial training, there was no difference in operant responding between the two replications (see Results: *Instrumental contingency versus random light delivery*). All other procedural details remained identical to Experiment 1.

### Procedure in Exp.3: Time course of response decrement and recovery (n = 7)

In the third experiment, mice from the first Y–maze experiment (after a 1-week break) were retrained with green light for 6 days under fixed-ratio and variable-ratio (VR) schedules. Animals were then tested for 7 days under FR5 with a dim red LED light to reveal the time course of response decrement. An additional test session with red light was conducted by a different experimenter (X.C.) after 7 days of red-light sessions conducted by S.K.E.T.; note that these mice were handled and tested solely by S.K.E.T. prior to the test session run by X.C.. The aim was to see if the attenuated operant response would recover when there were non-specific changes in the *extra*-experimental context—cues that animals might incidentally encode despite not being part of the behavioural task requirement (Hurst & West, 2010; Sorge et al., 2014; Sawangjit et al., 2022; Sakai et al., 2025). These sessions were conducted with a single lever in one arm of the Y–maze (**Figure 3*a***).

### Procedure in Exp.4: Interaction between food and light reinforcers (n = 8)

Animals’ body weights were recorded twice per 24 h at ZT02 and ZT14 for 7 days before the start of the experiment as well as 12 days during the experiment. In the acquisition phase (**Figure 4**), mice were maintained at 85–90% of their free-feeding body weights with ≤1.5 g food placed in their cages at ZT02 and ZT14 for 8 days. They were trained with a single lever in the operant chamber to work for a caloric and palatable food reinforcer under FR1 (sessions 1–6), FR3 (sessions 7 and 8), and FR5 (sessions 9–16). In sessions 13 and 14, food and 0.5-s green light onset occurred simultaneously (i.e., not forward *light* → *food* pairings that are optimal for establishing excitatory Pavlovian associations; Wheeler et al., 2008), to examine the unconditioned effect of light on steady-state appetitive responding when animals were in a food-restricted state. After session 16, mice were fed *ad libitum* until the end of the experiment. They were given test sessions with food reinforcers alone *versus* sessions with food delivered in conjunction with 2.5-s green light under FR5, to examine the effect of light when appetitive responding was not driven by caloric hunger but by the palatability of the food reinforcer. Food delivery and light onset occurred simultaneously, and the light reinforcer lasted for 2.5 s, during which further responding on the lever was recorded but not reinforced. Any potentiation of appetitive responding by light would not be driven by any Pavlovian-conditioned incentive due to a lack of forward *light* → *food* pairings (cf. Wheeler et al., 2008) before the test. The order of the two types of test sessions was summarised in **Supplementary Table 1*b***.

### Data analyses

Mean values ± standard errors of the mean (and medians in some cases) were reported in the main text and figures. Parametric analyses (*α* = 0.05, unless otherwise specified) were conducted in SPSS (version 29, IBM) and R.

In all experiments, primary analyses were within-subjects and split-plot analyses of variance (ANOVAs). We conducted higher-order ANOVAs to examine whether acquisition (indicated by the main effect of Session) and discrimination (indicated by the main effect of Lever) differed between test environments *via* relevant interaction terms (e.g., Lever × Setup, Session × Setup, and Session × Lever × Setup). This is because separate ANOVAs may lead to erroneous inferences, concluding that effects in the two setups differ simply because one is statistically significant and the other is not (Nieuwenhuis et al., 2011). Within-subjects factors in higher-order ANOVAs included training and test sessions, lever types (*L*+ *versus L*−), preexposure conditions (darkness *versus* green light), phases (day: ZT01–04 *versus* night: ZT13–16), and reinforcer types (food alone *versus* food and light); whereas between-subjects factors included experimental setups (operant chamber *versus* Y–maze), reinforcement contingency groups (Reinforced *versus* Yoked Control), and replications. For main and interaction effects involving within-subjects factors with more than two levels, Greenhouse–Geisser corrections were applied to the degrees of freedom (*df*) whenever the assumption of sphericity was violated. *F* values with adjusted *df* were denoted as *F_Ɛ_* in the main text. The primary dependent measure was operant responding, expressed as lever presses per 30 min or response rate per second; in some cases, operant responding was normalised to a reference condition to visualise within-subjects changes across mice (**Figures 1*e***, **3*b***–***g***, and **4*d***). Other dependent measures included the distance between the animal’s head and LED location, indicating the level of engagement with the goal of the task (**Figure 2*b***); and the total distance travelled, indicating the level of general activity. Distributions of inter-response times (IRTs), defined as the interval between each successive lever press, was visualised as heatmaps in Poincaré plots (**Supplementary Figure 3**).

In Experiments 3 and 4 where there were orderly changes in operant responding across sessions and individuals, a 3-paramater sigmoid function, *y* = *a***e**^−*x*^/[1+**e**^−*b*(*x*−*c*)^] or *y* = *a*/[1+**e**^−*b*(*x*−*c*)^], was fitted onto every individual response curve using nonlinear least-squares estimates obtained from the nls function in R (**Figures 3*c***,***f*** and **4*b***).

In Experiment 4, to examine the relative contributions of reinforcer summation (*food* and *light*), time of day (*night*), and body weight to appetitive responding, linear mixed-effects models were conducted using the `lmer` function (Bates et al., 2015), with individual mice as the random effect and lever pressing as the response variable. The importance of each fixed effect was assessed by examining the change in deviance (−2 × *maximum log-likelihood*) from comparing models with and without the fixed effect of interest. A significant drop in deviance, which was determined by the likelihood ratio *χ*^2^, indicated that incorporating the fixed effect in the model explained more variance in the response variable than removing it (Bates et al., 2015). The 95% confidence intervals of the fixed effects (i.e., slopes) were plotted in **Figures 4*e***,***f***.

Home cage PIR activity data with a temporal resolution of 10 s were smoothed with a 1-h moving average across the entire recording period. To visualise daily activity patterns, data were doubled-plotted in actograms (**Figure 4*a*** and **Supplementary Figure 4**). An activity midpoint was defined as the time point at which total activity in the preceding 8 h and total activity in the subsequent 8 h were equivalent. Specifically, for every time bin (*i*) the difference (Δ*_i_*) between preceding 8-h activity and subsequent 8-h activity was determined, and the product of Δ*_i_* and Δ*_i_*_−1_ was calculated. The time points at which Δ*_i_*Δ*_i_*_−1_ < 0 and Δ*_i_*_−1_ < 0 were defined as midpoints (Tam et al., 2021).

### Ethics

All behavioural procedures in this study were approved by the Duke Kushan University Institutional Animal Care and Use Committee (approved project no.: SKT-001) and conducted at the Duke Kunshan University—The First People’s Hospital of Kunshan Joint Brain Sciences Laboratory in accordance with the standard of the Association for Assessment and Accreditation of Laboratory Animal Care International.

## Results

### Exp.1: Two-choice discrimination

In the first experiment, we show that monochromatic light is an effective primary reinforcer that guides operant choice. Light self-administration training was conducted in the standard mouse operant chamber and Y–maze; the latter setup had a more complex spatial configuration than the former standard setup (**Figure 1*a***), which could affect operant choice. In both setups, there were two response levers, one was assigned as the correct lever (*L*+) and the other one as the incorrect lever (*L*−); positions of *L*+ and *L*− were counterbalanced across mice. Lever pressing on *L*+ with a force ≥2.5 g was reinforced; FR1 and FR3 were programmed in sessions 1–6 and sessions 7–12, respectively. Responding to *L*− did not have any consequence. The sensory reinforcer for *L*+ was presentation of monochromatic green LED light (*λ*_max_ = 535 nm) for 4 s. Green light was used because our previous study reported that light-induced anxiogenic response was lower under green light than isoquantal blue light (Pilorz et al., 2016), and a recent study reported the beneficial effect of green light on alleviating nociceptive pain in mice (Tang et al., 2022). Our animals were fed *ad libitum* in their home cages, so that lever responding was not driven by caloric hunger. Apart from this, our procedure was essentially identical to traditional operant paradigms with food and drug reinforcers. Unlike operant sensation seeking studies (Olsen & Winder, 2009; Gancarz et al., 2023), our sensory reinforcer remained unchanged in all sessions in the first experiment, so that responding in later sessions was unlikely to be driven solely by novelty of the light cue (as its novelty should *decrease* rather than increase as training progressed).

We found that monochromatic green light strengthened lever responding and guided operant choice in mice (**Figure 1*b***), indicating its reinforcing property. In cohorts *Operant*.*C1* and *Ymaze*.*C1* (*n* = 12; 1 non-responder excluded), responding on both levers increased across sessions [main effect of Session: *F*(1,11) = 34.93, *p* = 0.00017], and *L*+ responding was generally higher than *L*− responding [40.76 ± 9.40 *versus* 29.55 ± 6.64 presses; main effect of Lever: *F*(1,11) = 4.61, *p* = 0.055]. In a separate cohort of mice (*Operant*.*C2 n* = 8), we replicated the lever discrimination effect under FR5 in addition to FR1 and FR3 [main effect of Lever: *F*(1,7) = 5.98, *p* = 0.044; main effect of FR Schedule: *F*(2,14) = 3.51, *p* = 0.058]. FR Schedule × Lever interaction: *F*(2,14) = 2.59, *p* = 0.11]. Distributions of individual lever discrimination performance (i.e., the difference between *L*+ and *L*−) in cohorts *Operant*.*C1*, *Operant*.*C2*, and *Ymaze*.*C1* are depicted in **Supplementary Figure 1*a***; after outlier exclusion, the overall lever discrimination effect remained significant. Lever discrimination in the operant chamber and Y–maze was statistically indistinguishable despite the difference in spatial complexity of the environment [Lever × Setup interaction: *F*(1,19) = 1.09, *p* = 0.31; Session × Lever × Setup interaction: Greenhouse–Geisser corrected *F_Ɛ_*(3,52) = 0.78, *p* = 0.50]; separate Session × Lever ANOVAs for the two setups are shown in **Supplementary Table 2**. However, linear mixed-effects models revealed individual differences in learning rates—approximated by the slope of response change over training sessions—in both training environments. Between-subjects variance in learning rates was larger when training was conducted in the operant chamber than in the Y–maze (test of equality of variances: *F* = 4.09, *p* = 0.0078; **Supplementary Figure 2*b***), suggesting that individual learning trajectories are affected differently under different training environments.

### Exp.1: Satiation test

In most operant conditioning paradigms, lever pressing for food depends on whether rodents are hungry or satiated (Dickinson & Balleine, 1994). If light can serve as a reinforcer in a manner similar to food, light self-administration should be **“**satiable**”** to some extent by light preexposure. Moreover, satiation effects are more likely to be revealed if we take into consideration individual variability in operant behaviour, which is often reported in rodent operant studies (e.g., Dalley et al., 2007; Everitt et al., 2008; Ahmed, 2010; Pierce-Messick & Corbit, 2025). Thus, we subsequently conducted a subgroup analysis to examine satiation effects in High-responding *versus* Low-responding mice based on a median split.

To examine if light self-administration is sensitive to reinforcer satiation, we conducted test sessions after 30-min preexposure to darkness (the control condition) *versus* continuous green light in the operant chamber (*n* = 13) or in the start arm of the Y–maze (*n* = 7); levers were retracted during the preexposure period. After 30-min dark exposure, mice continued to respond more on *L*+ than *L*− [main effect of Lever: *F*(1,18) = 5.27, *p* = 0.034; Lever × Setup interaction: *F*(1,18) = 0.035, *p* = 0.85], but there was no difference between dark *versus* green light exposure [*L*+: *F*(1,19) = 2.96, *p* = 0.10; *L*−: *F*(1,19) = 1.17, *p* = 0.29]. Nevertheless, any satiation effect is less likely to be observed in animals with low levels of responding due to a floor effect. As such, we divided animals into two subgroups based on the median *L*+ response in the dark condition (*n* = 10 high *versus* low responders); the two subgroups showed comparable lever discrimination performance in the last two training sessions (main effect of Lever: *p* = 0.003; Lever × Responder Subgroup interaction: *p* = 0.37). Among high responders (**Figure 1*f***), 30-min light preexposure satiated *L*+ responding by 13% ± 5% (mean presses of 97.91 ± 19.88 *versus* 85.89 ± 16.72; *p* = 0.012) but no significant change was found on *L*− (mean presses of 34.53 ± 6.79 *versus* 48.10 ± 14.58; *p* = 0.077). No noticeable effect was found among low responders (*L*+: *p* = 0.88; *L*−: *p* = 0.78; **Figure 1*g***). These differential effects gave rise to a 3-way Preexposure Condition × Lever × Responder Subgroup interaction [*F*(1,18) = 6.66, *p* = 0.019]. Distributions of individual satiation effects (i.e., the difference between dark and green light preexposure) are depicted in **Supplementary Figure 1*b***; after outlier exclusion, the satiation effect on *L*+ in High-responders remained significant. However, the satiation effect was modest (∼13% reduction) and high responders were generally undeterred from self-administering light despite preexposure equivalent to hundreds of light presentations.

### Exp.2: Instrumental contingency versus random light delivery

Instead of serving as a primary reinforcer, light could have simply increased general responsiveness by changing the level of arousal or the vigilance state of the animal (Tam et al., 2020). To rule out these non-specific effects, a naïve cohort of mice (*n* = 21) was trained in one arm of the Y–maze with a single lever. Animals were assigned to two subgroups. In the Reinforced group (cohort *Ymaze*.*C2 n* = 11), there was *response* → *reinforcer* contingency such that the light reinforcer would be delivered whenever the FR response requirement was met. By contrast, no such instrumental contingency was programmed in the Yoked Control group, and the lever was permanently inactive (*n* = 10). Mice from Reinforced and Yoked Control groups were paired and run simultaneously in the left *versus* right arms of the Y–maze. Crucially, light delivery in the Control arm was **“**yoked**”** to the Reinforced arm, so that the control mouse received the same number of reinforcers at the same time as the experimental mouse during each session. All other procedural details remained identical to the first experiment.

The second experiment confirmed the role of monochromatic green light as a primary reinforcer in the operant task, rather than simply causing an increase in responsiveness in a non-specific manner. During baseline sessions when the Y–maze was kept in darkness (**Figure 1*c***), there was no difference in lever responding between subgroups [main effect of Subgroup: *F*(1,7) = 0.52, *p* = 0.50; Subgroup × Session: Greenhouse–Geisser corrected *F_Ɛ_*(2,12) = 1.21, *p* = 0.33]. From session 6 onwards, responding in the Reinforced group increased across sessions and reached a similar level as *L*+ responding in the first experiment. By contrast, responding in the Yoked Control group remained at the baseline level despite receiving equivalent light exposure as the Reinforced group [mean lever presses in Reinforced *versus* Yoked Control groups = 50.51 ± 10.97 *versus* 15.78 ± 4.05 presses; main effect of Group: *F*(1,17) = 6.70, *p* = 0.019; Group × Session interaction: Greenhouse–Geisser corrected *F_Ɛ_*(3,44) = 3.03, *p* = 0.046]; the latter interaction term was statistically indistinguishable between two partial replications [Group × Session × Replication interaction: Greenhouse–Geisser corrected *F_Ɛ_*(3,44) = 1.06, *p* = 0.37], indicating consistency across replications; separate ANOVAs for the two replications are shown in **Supplementary Table 3**. Moreover, in Reinforced mice there was a main effect of Ratio Schedule [*F*(2,20) = 5.07, *p* = 0.017] due to a linear increase in responding across FR1, FR3, and FR5 [*F*(1,10) = 13.68, *p* = 0.0041], similar to the increase in responding under ratio schedule progression in appetitive paradigms (see Experiment 4 below); no linear trend was found in Yoked Control mice [*F*(1,9) = 1.22, *p* = 0.30].

Moment-by-moment analyses revealed some key response structures in Reinforced mice; these characteristics are broadly consistent with typical ratio-schedule induced operant behaviour, thus distinguishing goal-directed lever presses in Reinforced mice from non-specific lever contact in control mice. Inter-response times (IRTs) were exponentially distributed with tightly clustered IRT distributions indicative of response bursts in Reinforced mice, whereas control mouse showed broader IRT distributions due to sparse lever engagement in the absence of instrumental contingency (**Supplementary Figure 3**). The group difference in lever responding is similarly evident shortly before and after light onset in 9 out of 10 of the Reinforced–Yoked Control pairs (**Figure 2*a***; except the last pair in which the yoked control mouse responded more than the Reinforced mouse). Notably, Reinforced mice showed reduced lever responding shortly after light onset [mean response rates during pre-LED *versus* post-LED periods = 0.088 ± 0.010 *versus* 0.069 ± 0.008 presses s^−1^; main effect of Period: *F*(1,10) = 7.98, *p* = 0.018; **Figure 2*a***]. This is consistent with the post-reinforcement pause that is a hallmark of ratio-schedule responding (Rickard et al., 2009). A further analysis on the distance between the animal’s head and LED location indicated that the decrease in responding during the post-reinforcement period was, at least in part, due to mice moving away from the lever and orienting towards the LED (**Figure 2*b***). Thus, in Reinforced mice lever presses that led to light delivery were accompanied by orienting responses towards the goal location, resembling Pavlovian conditioned approach responses (Robinson & Flagel, 2009); no goal-tracking behaviour was observed in Yoked Control mice (**Figure 2*b***).

### Exp.1 and Exp.2: Day versus night differences

The finding that nocturnal mice can learn to self-administer light seems to be at odds with the fact that they show spontaneous avoidance (Milosavljevic et al., 2016), enhanced learned fear (Warthen et al., 2011), elevated cardiac response (Thompson et al., 2008), and hypothalamic–adrenal mediated stress response to light (Ishida et al., 2005). These effects are observed in the light phase but tend to be stronger at night (Ishida et al., 2005). Given that behavioural, physiological, and brain responses of laboratory mice are highly sensitive to light during the first few hours of the biological night but less so during the light phase (Foster et al., 2020), there could be a time-of-day difference in light self-administration. To investigate this, we compared operant responding in the final FR1, FR3, and FR5 sessions conducted at Zeitgeber time (ZT)01–04 and ZT13–16 (**Figure 1*d***). Mice showed reduced *L*+ responding at night [FR1: *F*(1,29) = 7.09, *p* = 0.013; FR3: *F*(1,29) = 9.11, *p* = 0.0053; FR5: *F*(1,17) = 5.91, *p* = 0.026], indicative of a reduction in the reinforcing property of light—in contrast to the increase in food self-administration at night (**Figure 4*c***)—suggesting that responding for light and responding for food are regulated differently. The time-of-day effect was lever specific, as it was more significant for *L*+ than *L*− [FR1: *F*(1,29) = 0.10, *p* = 0.75; FR3: *F*(1,29) = 2.42, *p* = 0.13; FR5: *F*(1,17) = 4.70, *p* = 0.045; Time of day × Lever interaction: *F*(1,28) = 10.37, *p* = 0.0032; Time of day × Lever × FR Schedule interaction: *F*(1,28) = 3.66, *p* = 0.066]. Distributions of individual time-of-day effects (i.e., the difference between day and night) are depicted in **Supplementary Figure 1*c***; after outlier exclusion, time-of-day effects on *L*+ responding under FR1, FR3, and FR5 remained significant. Interestingly, *L*+ responding at night could be reinstated by lengthening the duration of the light reinforcer [*F*(1,17) = 13.02, *p* = 0.002; **Figure 1*e***]. This indicates that a change in the physical property of the light cue may override the reduced reinforcing property of light at night, further indicating that, unlike previous studies that used light of longer durations (Ishida et al., 2005; Milosavljevic et al., 2016), the light cue in our paradigm is not *strictly* anxiogenic or stress inducing.

### Exp.3: Response decrement under red light

Well-established behaviour often takes time to modify (Hull, 1943). In standard extinction procedures (i.e., no reinforcers), operant responding acquired under FR schedules often persists for multiple extinction sessions before declining to the pre-training level in rodents (e.g., Shaw et al., 2004; Marini et al., 2025). But instead of using standard extinction procedures without reinforcement to modify the response, we examined if well-established responding could be weakened when the green LED light was replaced by a dim red LED light (*λ*_max_ = 630 nm; **Figure 3*a***), which emitted a comparable level of heat (**Figure 3*b***) but excited the mouse retinal photoreceptors (*S*-cone, melanopsin, rod, and *M*-cone) to a lesser extent (**Figure 3*c***). This allowed us to determine if operant responding could decrease with an attenuated photic signal (resulting in response decrement) or whether responding could be sustained by non-photic, LED-emitted thermal cues alone—which are known to be effective primary reinforcers in mice (Gordon, 1985; Gordon et al., 1998).

After the initial training in the Y–maze in Experiment 1, cohort *Ymaze*.*C1* (*n* = 7) were retrained with green light for 6 days under fixed-ratio and variable-ratio (VR) schedules in a counterbalanced order: FR1, FR3, FR5, VR5, VR5, and FR5. The overtrained operant response was stable with no difference between FR5 and VR5 [74.50 ± 10.60 *versus* 69.50 ± 7.40 presses, respectively; *F*(1,6) = 1.40, *p* = 0.28]. After confirming that operant responding was stabilised, mice were tested for 7 days with a dim red LED light to reveal the time course of response decrement. Red light continued to be delivered under FR5, so that experimental sessions were matched as closely as possible to retraining sessions in terms of non-photic factors (i.e., LED-generated heat). After switching to red light, operant responding remained at comparable levels for several days without any significant change, but gradually decreasing to ∼50% after a week (**Figure 3*e***,***h***). Response decrement was accompanied by a slight decrease in locomotor activity and a slight increase in proximity to the LED location on the last day (**Figure 3*d***,***g***). The gradual decline in light self-administration resembles the time course of extinction of food, sucrose, and drug self-administration under ratio schedules (FR1–FR5), typically taking about a week for responding to return to the pre-training level (Shaw et al., 2004; Keiflin et al., 2008; Marini et al., 2025). Although lever responding became weaker under red light in our experiment, it was partially recovered following an unexpected change in the experimenter on the probe session [*t*(6) = −3.87, *p* = 0.0082 (2-tailed); **Figures 3*f***,***i***]. This suggests response decrement is under some contextual control by *extra*-experimental cues such as handling (Hurst & West, 2010) or experimenter-related odours (Sorge et al., 2014).

### Exp.4: Interaction between light and food reinforcers

Finally, to systematically determine previously reported interactions between light and other primary reinforcers (Caggiula et al., 2002; Wolff & Saunders, 2024), we first trained a naïve cohort of mice (*n* = 8) in a standard food self-administration task with a single lever in the dark operant chamber to obtain a 20-mg cream-flavoured food pellet (**Figures 4*a***–***c***). These animals were food restricted during FR1, FR3, and FR5 training. After acquisition, animals were fed *ad libitum*; operant responding for food persisted when mice were not hungry [FR1 *versus* FR5 fed *ad libitum*: 52.43 ± 7.83 *versus* 110.25 ± 6.67 presses; *t*(7) = −6.65, *p* = 0.00029 (2-tailed)], albeit at 36% ± 4% of the steady state [FR5 food restricted *versus* fed *ad libitum*: *t*(7) = 15.64, *p* < 0.00001 (2-tailed)]. On test sessions, we compared lever responding to food reinforcers alone *versus* food reinforcers delivered simultaneously with the onset of 2.5-s green light under FR5. Light potentiated operant responding for food when mice were fed *ad libitum* (**Figure 4*d***,***f***) but not when food restricted (**Figure 4*e***), confirming the additive nature of the reinforcing effectiveness of light and a palatable reinforcer. The effect of light was not driven by any Pavlovian-conditioned incentive, because simultaneous *light*–*food* pairings are known to be suboptimal for establishing excitatory Pavlovian associations (Wheeler et al., 2008).

## Discussion

Under typical housing conditions, laboratory mice are nocturnal; they are often considered to be highly sensitive to environmental light (Foster et al., 2020) and show bright-light avoidance in classic tests of anxiety (Semo et al., 2010), including light/dark box, elevated–plus maze, and open field tests, as well as in the home cage when a dark receptacle is available (Steel et al., 2024). The fact that they are willing to work for light (each lever press requiring ≥2.5 g i.e., ∼10% of an average male C57BL/6 mouse’s body weight of 25 g) under non-physiologically deprived conditions has never been formally investigated.

### Light as a reward to establish habits: A summary

Key results from our four experiments can be summarised as follows: (*a*) A brief green-light cue that is presented immediately with a lever press can increase lever responding under traditional fixed-ratio schedules of reinforcement. Mice press more on the lever that is paired with green light than the lever that has no consequence (**Figure 1*b***, **Supplementary Figure 1*a***, and **Supplementary Table 2**). (*b*) Compared with dark preexposure, green-light preexposure results in reduced lever pressing on the reinforced lever among High-responding mice. However, this effect is modest as these mice continue to press the reinforced lever at a similar level (**Figure 1*f*** and **Supplementary Figure 1*b***). (*c*) The increase in lever responding requires learning about the relationship between lever presses and light cue delivery i.e., it requires *response* → *reinforcer* (instrumental) contingency; the same number of light cues presented at the same moments in time but in a random manner (i.e., zero contingency) does not support acquisition of lever responding (**Figure 1*c*** and **Supplementary Table 3**). Response characteristics of lever responding in the contingent (but not non-contingent) group conform to what we know about appetitively motivated behaviour under fixed-ratio schedules (**Figure 2** and **Supplementary Figure 2**). (*d*) Once lever responding is at a stable level, it is possible to reduce responding by replacing green light with red light (**Figures 3*d***,***e***,***g***,***h***)—which can be detected by mice but activates the mouse’s photoreceptors to a lesser extent than green light (**Figure 3*c***). The response decrement under red light further confirms that some behaviour is indeed acquired in the preceding phase with green light. (*e*) In mice that are well trained to lever-press to obtain food, the unexpected presentation of a brief green-light cue delivered in conjunction with the food pellet enhances lever responding (**Figure 4*d***). Lever responding is potentiated by light when mice are fed *ad libitum* (**Figure 4*f***) but not when food restricted (**Figure 4*e***). Collectively, our study demonstrates that response-contingent light can serve as a reward in mice.

### Light versus drug reinforcers: A comparison

Compared with other operant drug studies, the average asymptotic level of responding in our light-based paradigm is slightly higher than in recent rat operant drug studies under FR1. For example, in Seaman et al. (2022) after 7 days of operant self-administration training with methamphetamine and fentanyl, the mean asymptotic level of *L*+ responding was <20 lever presses in a 90-min session for both drugs; this is lower than what we observed under FR1 (our session duration was shorter than theirs). Similarly, in Marini et al. (2025) after 10 days of operant heroin self-administration training, the mean asymptotic level of *L*+ responding was ∼30 lever presses (session duration was not reported), again lower than in our **Figure 1*b***. However, lever discrimination effects in these operant drug studies are generally stronger than in our study. This suggests that, although light can sustain responding, it may be less effective than drug reinforcers in guiding behavioural choice.

### Non-specific effect: Alerting effects of light and LED-generated heat

Our results cannot be explained in terms of a single non-specific effect. For example, light exerts an alert-promoting effect on general activity in mice (Thompson et al., 2008), potentially increasing lever contact in a non-specific manner. However, this cannot explain the observed differences between contingent and non-contingent groups in Experiment 2, as both are exposed to the same number of light cues at the same moment during the training session. The contingent group showed a characteristic orienting response towards the goal after a rewarded lever press (**Figure 2*b***). This resembles goal tracking, a form of Pavlovian conditioned approach responses elicited by the lever serving as a conditioned stimulus (Robinson & Flagel, 2009). By contrast, the non-contingent group did not show any sign of goal tracking, suggesting that light exposure *per se* is not sufficient for any anticipatory response to be established. Thus, instrumental and Pavlovian processes may drive different components of light-seeking behaviour, neither of which can be explained in terms of alerting effects of light (as this should be equated in contingent and non-contingent groups). The current analysis of goal-tracking behaviour is exploratory; this will need to be confirmed in future studies with more complex experimental designs.

It is also possible that lever responding is sustained by LED-generated non-photic rather than photic cues, as thermal cues are effective primary reinforcers that guide behavioural choice in mice (Gordon, 1985; Gordon et al., 1998). For example, the ambient temperature in a typical housing condition for laboratory mice is ∼21°C; but when they are given the chance to select different levels of ambient temperature (*T_a_*) in a thermally graded environment, mice tend to choose *T_a_* > 27°C in the light phase and *T_a_* > 25°C at night (Gordon et al., 1998). By using a red LED light that activates the mouse’s photoreceptors to a lesser extent but generates a comparable level of heat (**Figure 3*b***), we demonstrate that lever responding cannot be sustained by thermal cues alone. While our experimental conditions demonstrate the reinforcing value of green light (and the relatively limited reinforcing value of dim red light), other wavelengths (e.g., UV) may also have similar reinforcing properties in mice (Fell et al., 2014); this remains to be investigated in future operant studies.

### State-dependent effects of light

Although our study confirms the reinforcing property of green light at a certain intensity, when light becomes too bright (>1000 lux) or when its duration becomes too long (e.g., in the minutes-to-hours range), its reinforcing value is counteracted by other factors such as light-induced stress (Ishida et al., 2005; Thompson et al., 2008), resulting in anxiogenic responses such as light avoidance in the light/dark box test (Semo et al., 2010; Milosavljevic et al., 2016; Wang et al., 2023). Thus, light can induce either approach or avoidance response (i.e., the effect on behaviour is bidirectional), depending on light intensity, test context, and the mouse’s vigilance state (Tam et al., 2020). The relationship between light intensity and its reinforcing value is likely to be complex and nonmonotonic, but it should generally follow an inverted-U i.e., Yerkes–Dodson-type function (Tam et al., 2025).

As light can exert a state-dependent effect on the mouse’s behaviour (Tam et al., 2020), the day–night difference in light self-administration can be interpreted in terms of approach–avoidance conflict (Gray, 1987; McNaughton, 2025) due to differential light sensitivity across the daily cycle (Foster et al., 2020). Specifically, according to the mouse’s phase-response curve (PRC) mice exhibit higher sensitivity to light during the early subjective night around dusk than during the early subjective day, which is labelled as the **“***dead zone***”** (Foster et al., 2020). Heightened sensitivity to nocturnal light may amplify its arousal effect and counteract its reinforcing value, leading to reduced light seeking (i.e., *avoidance* ≫ *approach*), whereas the approach response may dominate in the light phase (i.e., *approach* ≫ *avoidance*). Furthermore, mice encode stimulus durations in conditioning procedures (Gallistel & Gibbon, 2000) and update temporal information rapidly (Aggadi et al., 2025). Thus, a change in light duration that violates learned expectations may enhance attentional processing of the cue (Tang et al., 2023). This may transiently shift the balance towards approach behaviour, thereby reinstating lever pressing that is otherwise suppressed at night; this notion remains to be tested in future studies.

### Contextual influences: Training context and extra-experimental cues

Given that context can affect drug self-administration, we use two distinct environments to examine the influence of training context on light self-administration. Notably, context novelty can enhance acquisition of cocaine but not saline self-administration in rats (Caprioli et al., 2007), highlighting the role of context in drug-seeking habit formation. In our study, there is no group difference in acquisition or lever discrimination performance between the operant chamber and Y–maze (although the latter is more geometrically distinct from the mouse’s home cage). However, there is individual variability in learning rates (**Supplementary Figure 2**); in particular, between-subject variance in learning rates is slightly higher in the operant chamber than in the Y–maze, suggesting that although average performance is similar, training context may affect individual learning trajectories differently. Beyond training context, modulation can arise from *extra*-experimental context—cues that animals incidentally encode despite *not* being part of the behavioural task requirement (Welker & McAuley, 1978; Hurst & West, 2010; Sorge et al., 2014; Sawangjit et al., 2022; Sakai et al., 2025). The (partial) response recovery under red light following experimenter change (**Figures 3*f***,***i***) shows that operant response decrement in our paradigm is under contextual control, potentially by handling cues (Hurst & West, 2010) or experimenter-related odours (Sorge et al., 2014). This resembles the context specificity of extinction of food-seeking and drug-seeking habits: response decrement may involve formation of inhibitory *context*–*response* associations that interfere (retrospectively) with the old habit (Bouton, 1993, 2019, 2024), rather than weakening of the old habit. Together, our results provide some initial evidence that light self-administration can be modulated by nonspecific extra-experimental context as well as directly affected by training context.

### Bridging the gap between animal models and human studies

The light self-administration paradigm in mice is relevant to behavioural addiction, in particular, digital technology-based disorders such as smartphone use and gaming disorder (Tam et al., 2025). Several recent studies have reported that coloured light emitted from digital devices can influence daily usage. By applying grayscale filters to smartphone displays, screen time can be reduced by ∼20–50 min per day in young adults (Holte & Ferraro, 2020; Dekker & Baumgartner, 2023; Holte et al., 2023; Wickord & Quaiser-Pohl, 2023; Zimmermann & Sobolev, 2023). These results confirm the notion that coloured light possesses some reinforcing values that exacerbate maladaptive engagement with operandum-*like* devices in humans (e.g., fruit machines; Griffiths, 1993). Light is also known to have some therapeutic values in treating certain psychiatric disorders (see e.g., Menegaz de Almeida et al., 2025, for a recent meta-analysis on the efficacy of bright light therapy over red light exposure for alleviating depressive symptoms); but the mechanisms underlying its effectiveness remains to be investigated (Huang et al., 2024). In relation to behavioural addiction, a recent study (Li et al., 2026) has reported the effectiveness of light therapy (*versus* placebo) in reducing symptom severity among participants with symptoms of Internet gaming disorder (IGD). Functional magnetic resonance imaging (fMRI) analysis has revealed functional connectivity changes between the orbitofrontal cortex (OFC) and habenula and between the OFC and ventral tegmental area (VTA) after light therapy that correlate with changes in cue-induced craving symptoms (Li et al., 2026). Thus, light can modify human brain networks that are known to contribute to compulsive and addictive behaviours (Koob & Volkow, 2016). The light-based operant paradigm allows us to examine how brain reward circuits are (re)shaped by light contingencies; it also permits initial screening of the effectiveness of light-based *versus* novel pharmacological interventions for (behavioural) addictions in experimental animals before being translated into human studies (cf. the recent study by Soler-Cedeno et al., 2025, screening a novel dopamine transporter inhibitor).

### Validation of a new operant paradigm

Here we validate a light-based operant paradigm using traditional ratio schedules of reinforcement; no palatable food or drug reinforcer is needed to motivate the mouse to self-administer light. This behaviour is not driven by the urge to seek novelty or surprise (unlike Olsen & Winder, 2009; Gancarz et al., 2023), as the light reinforcer remains unchanged during training in Experiment 1 (for cohorts *Operant*.*C1*, *Operant*.*C2*, and *Ymaze*.*C1*).

Importantly, the light self-administration paradigm induces persistent behaviour that cannot be fully **“**satiated**”** (in High-responding mice) despite light preexposure equivalent to hundreds of light-cue presentations—a phenotype potentially relevant to the study of human behavioural addiction. While our findings demonstrate that light can support persistent responding in some individuals, it remains to be determined whether the behaviour observed here would meet stricter criteria for habitual responding. Future studies employing outcome-devaluation or contingency-degradation procedures are required to test whether responding becomes insensitive to outcome value or *action*–*outcome* contingency (Dickinson & Balleine, 1994; Balleine & Dickinson, 1998; Yin & Knowlton, 2006).

Over the course of our study, we have noticed an animal-welfare related advantage of the light-based paradigm over appetitive operant paradigms: the latter requires restricted feeding, which disrupts the mouse’s natural rhythm (**Supplementary Figures 4*a***,***b***). This unintended consequence can be circumvented by operant training with brief light cues that does not require food restriction, thus maintaining the mouse’s nocturnality during the experiment (**Supplementary Figures 4*c***,***d***).

## Conclusions

The light self-administration paradigm expands the class of non-food, non-drug primary reinforcers (thermal cues, running wheel, novel objects, and conspecifics of different sexes) that are deemed effective in mice (Gordon et al., 1998; Bevins & Basher, 2005; Ramsey et al., 2023; Nishitani et al., 2025), allowing the assessment of maladaptive behaviour driven neither by physiological needs nor the urge to seek novelty. Several key questions remain to be investigated; these are listed in **Supplementary Table 4**. Our study represents the initial step for developing more sophisticated experimental models for disorders characterised by compulsions including digital technology-based disorders (Casile et al., 2025; Tam et al., 2025). The validated paradigm can help to understand how interactions between visual and reward circuits give rise to the reinforcing property of light and light-motivated behaviour (Tam et al., 2025). It may help to reveal common (i.e., transdiagnostic) mechanisms that are shared across multiple compulsive and addictive behaviours (Klugah-Brown et al., 2021; Robbins et al., 2024).

---

**Supplementary Figure 1.**
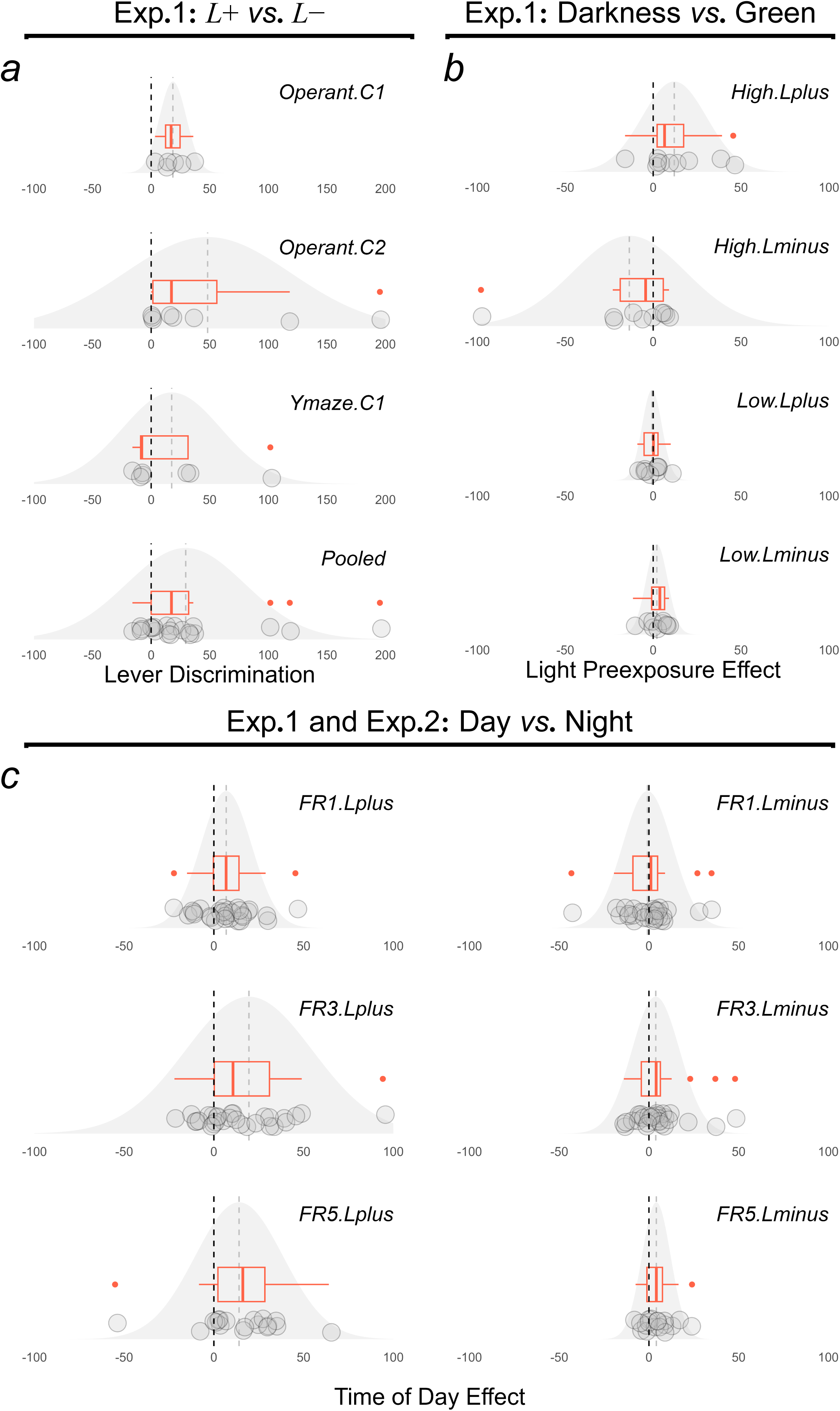
| Distributions of lever discrimination performance, light preexposure (satiation) effects, and time-of-day effects in Experiments 1 and 2. (a) Data were from the last day of operant training under FR3 (session 12) in Experiment 1. There were *n* = 6, *n* = 8, and *n* = 7 mice trained in cohorts *Operant*.*C1*, *Operant*.*C2*, and *Ymaze*.*C1*, respectively, as in **Supplementary Table 2**. All trained mice were included in this panel, including the mouse that was classified as a non-responder in Figure 1a. *Black* dashed lines at zero indicate no discrimination performance (i.e., *L*+ presses ≈ *L*− presses). *Grey* dashed lines represent the mean difference of *L*+ and *L*− responding. Outliers with *L*+ responding >120 presses on the last day of FR3 training are indicated by ● above *grey* data points. In the *Pooled* panel (panel ***a***, *bottom*), both parametric and non-parametric analyses showed that the lever discrimination effect was insensitive to outliers: ANOVA with outliers, *F*(1,20) = 7.14, *p* = 0.015, partial *η*^2^ = 0.26; ANOVA without outliers, *F*(1,17) = 8.33, *p* = 0.010, partial *η*^2^ = 0.33; non-parametric test with outliers, Friedman *χ*^2^ = 6.37, *p* = 0.012 (2-tailed); and non-parametric test without outliers, Friedman *χ*^2^ = 4.00, *p* = 0.046 (2-tailed). (***b***) Data were from light preexposure test sessions in Experiment 1. Mice were divided into two subgroups (High *versus* Low responders) based on the median *L*+ response in the dark preexposure condition. *Black* dashed lines at zero indicate no light preexposure effect (i.e., lever presses remained the same after preexposure to darkness and green light). *Grey* dashed lines represent the mean difference between darkness and green light conditions. In the High-responder subgroup (panel ***b***, *top*), there was a satiation effect of green light on *L*+ responding. The leave-one-out (jackknife) approach was used to examine the influence of any outlier. Specifically, each mouse in the High-responder subgroup was omitted sequentially and the main effect of light preexposure was examined with *n* − 1 cases; all (but one) *p* values from the ANOVA remain significant: *F*s ≥ 5.21, *p*s ≤ 0.056. (***c***) Data were from main Figures 1d,***e*** but replotted to show the difference in lever presses during sessions conducted in the light phase [Zeitgeber time (ZT)01–04] *versus* at night (ZT13–16). *Black* dashed lines at zero indicate no time-of-day effect (i.e., lever presses did not differ between day and night). *Grey* dashed lines represent mean differences, which were greater than zero for *L*+ (i.e., *Day* > *Night*; panels ***c***, *left*) and closer to zero for *L*− (i.e., *Day* ≈ *Night*; panels ***c***, *right*). Time-of-day effects on *L*+ responding were insensitive to outliers, as effects remained significant after outlier exclusion: FR1, *F*(1,23) = 6.88, *p* = 0.023; FR3, *F*(1,23) = 12.44, *p* = 0.0018; and FR5, *F*(1,14) = 14.36, *p* = 0.0020. These effects do not interact with Setup (operant chamber *versus* Y–maze): FR1, *F*(1,23) = 0.13, *p* = 0.72; FR3, *F*(1,23) = 0.01, *p* = 0.93; and FR5, *F*(1,14) = 0.98, *p* = 0.34. For all panels, between-subject variance is visualised as the spread of the fitted normal distribution in *grey*. In each boxplot (***red***), the thick central line represents the median difference, and the width of the box indicates the interquartile range (IQR, which is *Q*_1_–*Q*_3_). Data points that are 1.5 × IQR < *Q*_1_ or 1.5 × IQR > *Q*_3_ are considered as outliers (●). **“**Whiskers**”** of the box extend from the IQR to minimum and maximum values excluding identified outliers. Boxplots are generated by the `geom_boxplot` function in R.

**Supplementary Figure 2.**
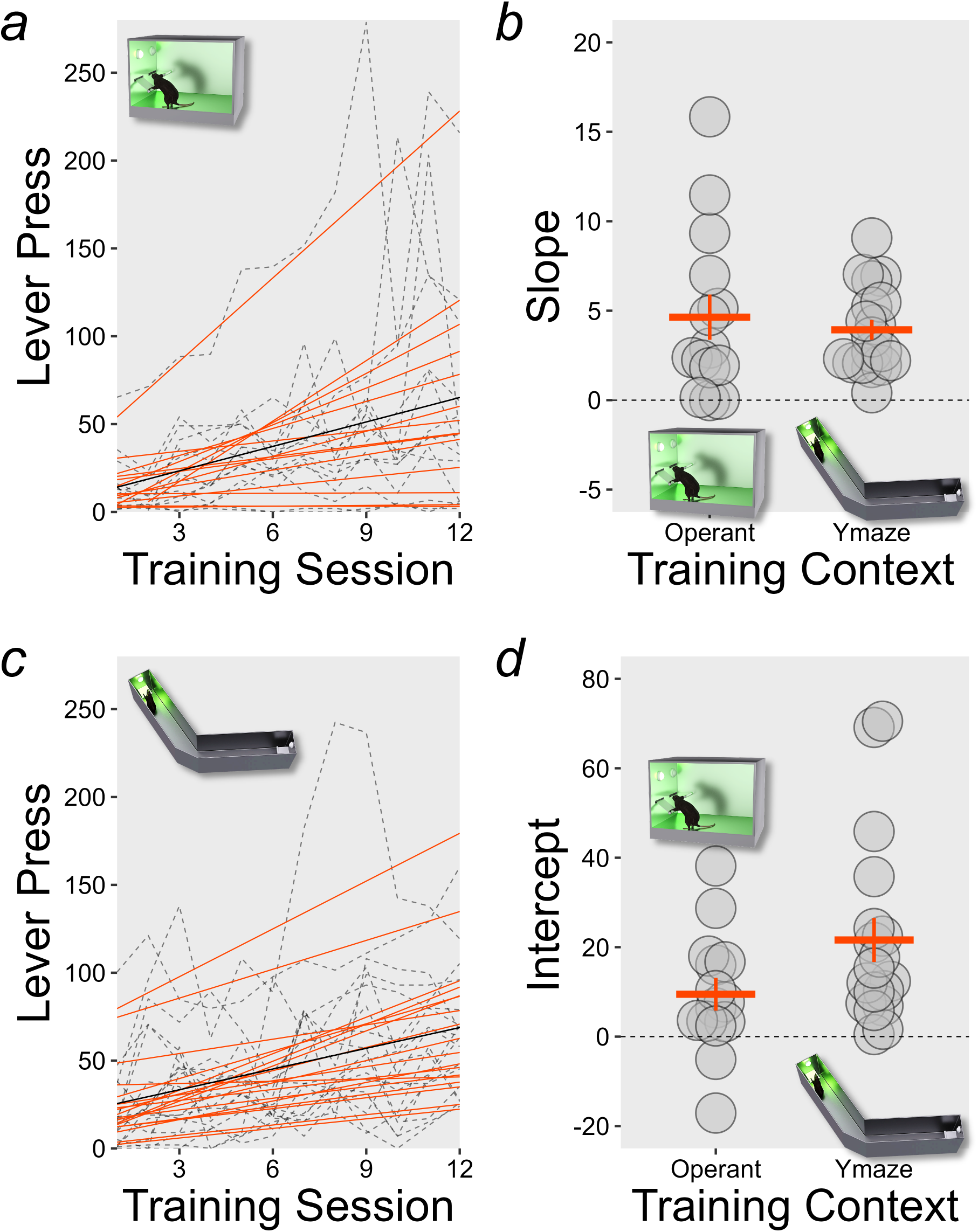
| Individual variation in learning to self-administer light. (***a***, ***c***) Lever responding on *L*+ in the first 12 training sessions (*grey dashed lines*) was fitted with a linear model (*black lines*) and a linear mixed-effects model (*red lines*) with random slope and intercept (Bates et al., 2015). Fitting was done separately for cohorts trained in the operant chamber, *Operant*.*C1* (*O*.*C1*) and *Operant*.*C2* (*O*.*C2*) in panel ***a***, *versus* cohorts trained in the Y–maze, *Ymaze*.*C1* (*Y*.*C2*) and *Ymaze*.*C2* (*Y*.*C2*) in panel ***c***. Individual differences in learning rates can be captured by the random effect in the linear mixed-effects model, but not the simple linear model with group-level slope and intercept estimates. In both setups, there were positive fixed effects of Training Session (operant chamber: *χ*^2^ = 8.25, *p* = 0.0041; Y–maze: *χ*^2^ = 16.90, *p* < 0.0001), confirming the acquisition effects from ANOVAs. (***b***, ***d***) Mean slope estimates (***red***) in panel ***b*** were not different between setups [*F*(1,30) = 0.31, *p* = 0.58], but between-subjects variance was larger in the operant chamber than in the Y–maze (test of equality of variances: *F* = 4.09, *p* = 0.0078). With respect to intercept estimates in panel ***d***, between-subjects variance was comparable between setups (test of equality of variances: *F* = 0.43, *p* = 0.13). However, there was a nonsignificant trend of higher intercepts in the Y–maze [*F*(1,30) = 3.51, *p* = 0.071], reflecting slightly more responding in initial training sessions.

**Supplementary Figure 3.**
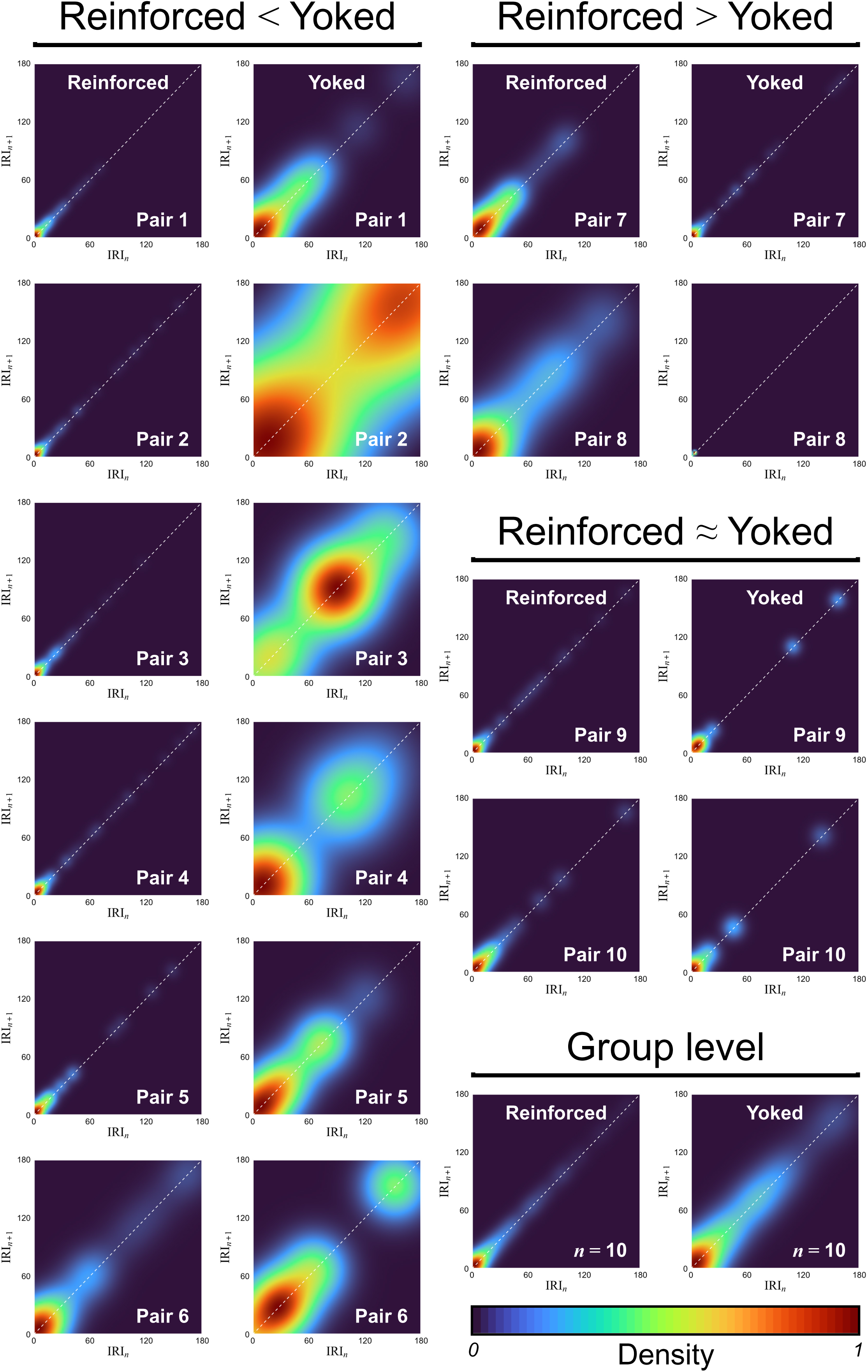
| Inter-response times (IRTs) under FR5. For each mouse, IRT variability was visualised in Poincaré plots as density heatmaps using `stat_density_2d` in R. Each density plot shows the correlation between successive inter-response intervals [(IRI_1_,IRI_2_), (IRI_2_,IRI_3_), (IRI_3_,IRI_4_),**…**(IRI*_n_*,IRI*_n_*_+1_)**…**] using data from the final session in Experiment 2. The diagonal dashed line is the reference line IRI*_n_* = IRI*_n_*_+1_. In 6 out of 10 Reinforced–Yoked Control pairs (pairs 1–6), the control mouse showed a higher IRT variability (i.e., a wider density distribution) than the Reinforced counterpart, indicating less consistency in operant responding and therefore less engagement with the operandum. The remaining pairs either showed the opposite pattern (in pairs 7 and 8) or no obvious difference (in pairs 9 and 10). Group-level density heatmaps were also generated by cojoining response records of all mice from end to end; note that one Reinforced mouse is not included, as it did not have a Yoked Control counterpart (see Methods: *Animals and housing condition*). Group-level density plots retained the overall pattern of a higher IRT variability in the Yoked Control group than the Reinforced group, broadly consistent with the observations in the majority (6 out of 10) of Reinforced–Yoked Control pairs.

**Supplementary Figure 4.**
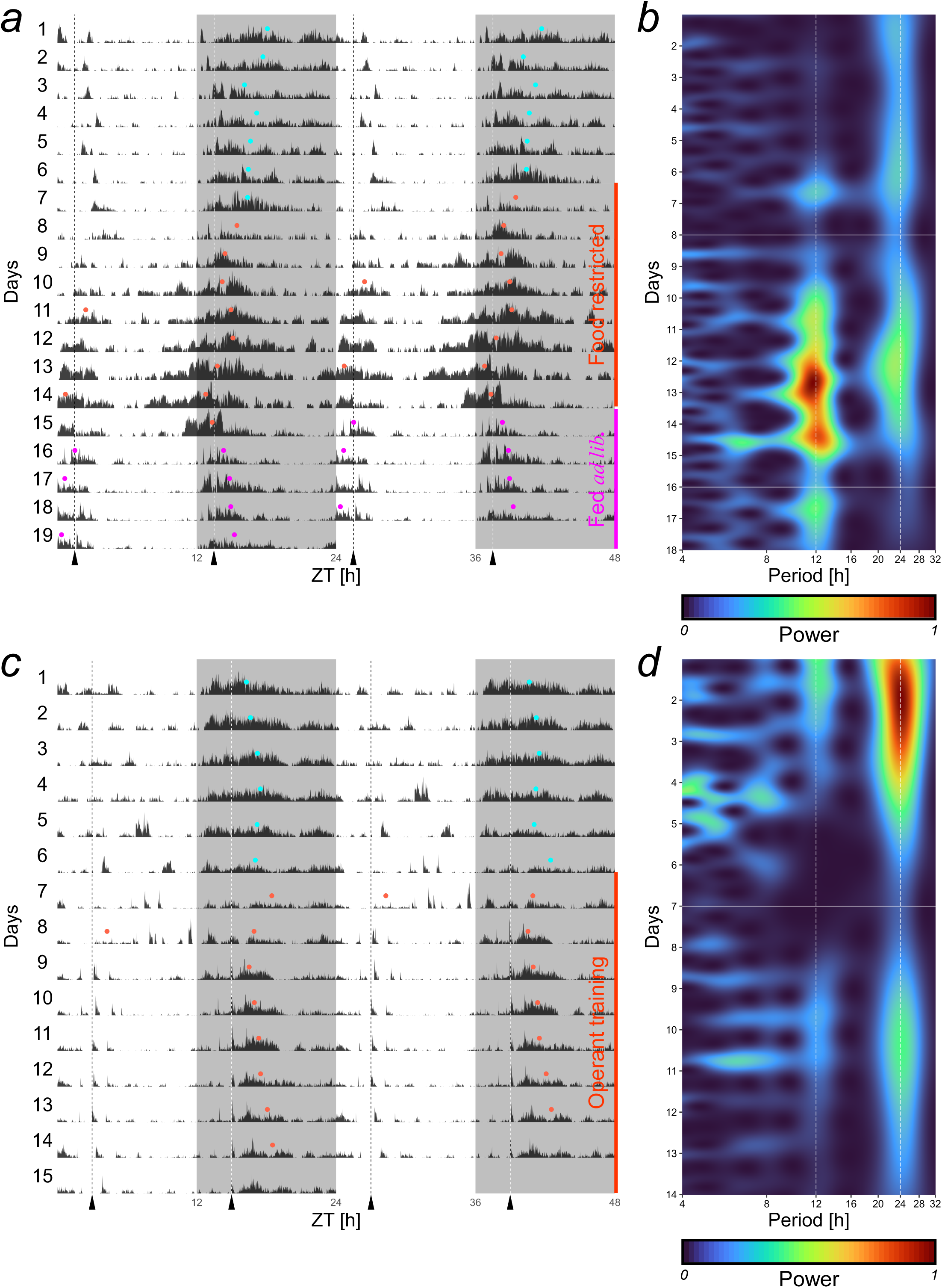
| Training for food but not light disrupts nocturnality. (***a***) The double-plotted actogram (data pooled across mice) shows home cage activity from Experiment 4 as described in Figure 4a. Before the experiment commenced, there was only one activity midpoint per day. By contrast, there were two activity midpoints on some days during operant training, indicating that it disrupted nocturnality (van der Vinne et al., 2014). Midpoints were close to feeding times represented by vertical dashed lines (*black* and *white*). (***b***) Wavelet analysis of the time series from panel ***a*** (using `WaveletComp`; Rösch & Schmidbauer, 2019) revealed that activity rhythm oscillated at 24 h before the experiment. However, non-24-h oscillations centred at 12 h dominated during restricted feeding on days 8–15, persisting beyond day 15 when mice were fed *ad libitum*. This indicates entrainment of activity rhythm to the feeding cycle (i.e., food earned from the operant task or given by the experimenter) but not to the light:dark cycle. (***c***) The double-plotted actogram shows home cage activity before and during light self-administration training in Experiment 1; data were pooled across 6 mice under home cage activity monitoring. Vertical dashed lines indicate training times. There was only one activity midpoint per day throughout the experiment (except day 8). (***d***) Wavelet analysis of the time series from panel ***c*** did not show any noticeable increase in power at the 12-h period.

**Supplementary Table 1:**
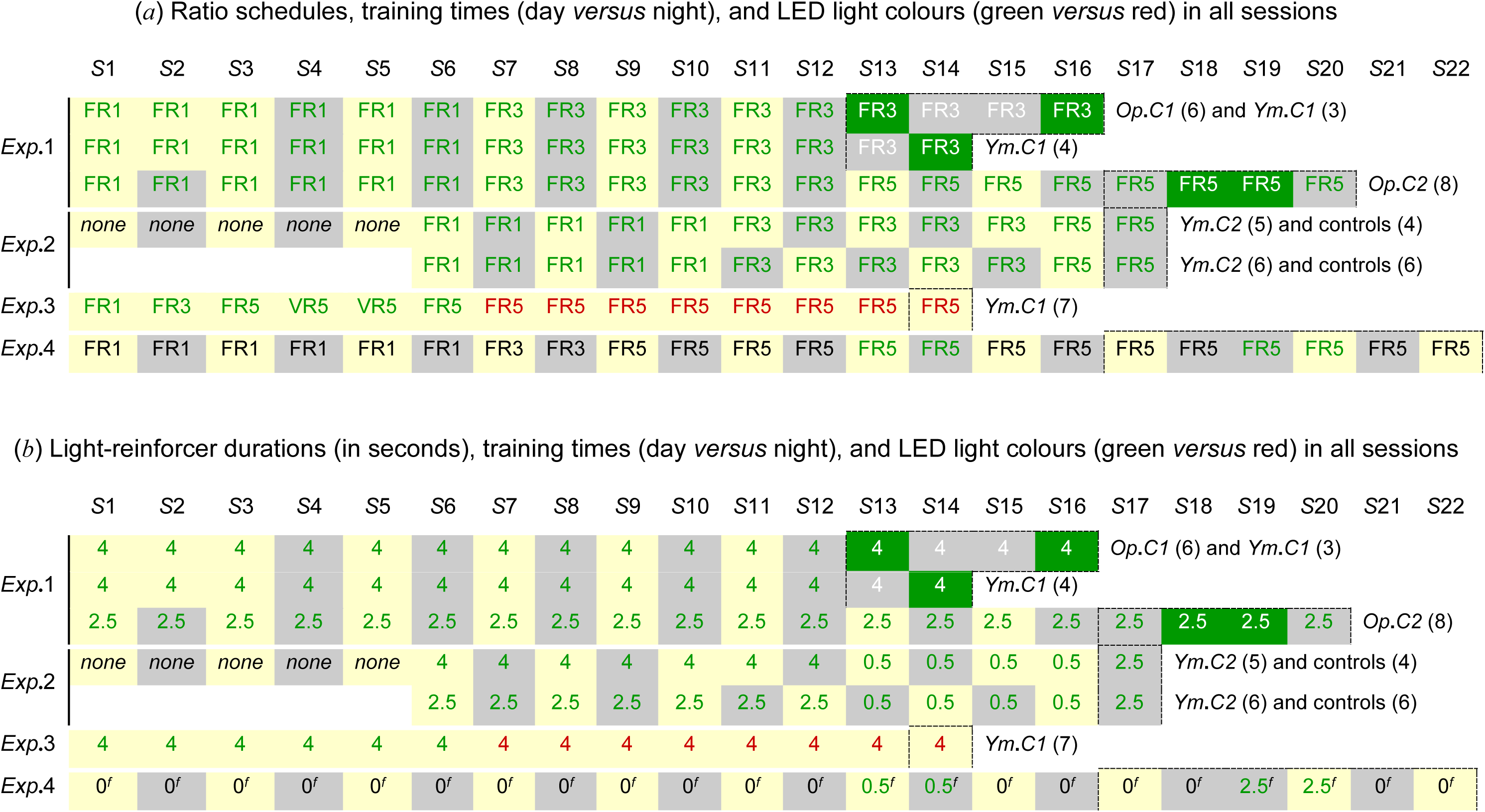
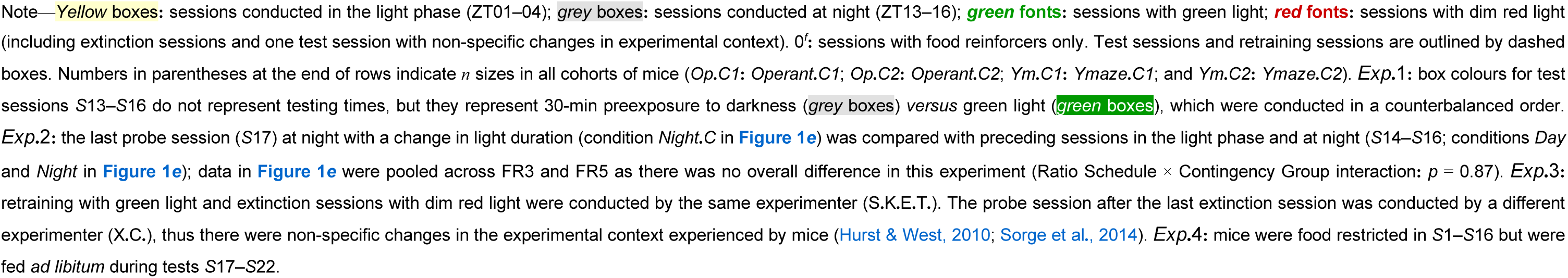

**Supplementary Table 2:**

| Cohort | Main effect of<br>Session (*acquisition*) | Main effect of<br>Lever (*discrimination*) | Session × Lever<br>interaction |
| --- | --- | --- | --- |
| *Operant.C1*<br>$n = 6$ | $F(11,55) = 3.56$<br>$p = 0.00082^{**}$<br>partial $\eta^2 = 0.42$ | $F(1,5) = 7.45$<br>$p = 0.041^*$<br>partial $\eta^2 = 0.60$ | $F(11,55) = 1.96$<br>$p = 0.051^*$<br>partial $\eta^2 = 0.28$ |
| *Operant.C2*<br>$n = 8$ | $F(11,77) = 3.59$<br>$p = 0.00041^{**}$<br>partial $\eta^2 = 0.34$ | $F(1,7) = 5.91$<br>$p = 0.045^*$<br>partial $\eta^2 = 0.46$ | $F(11,77) = 2.76$<br>$p = 0.0045^{**}$<br>partial $\eta^2 = 0.28$ |
| *Ymaze.C1*<br>$n = 7^\dagger$ | $F(11,66) = 10.65$<br>$p < 0.0001^{**}$<br>partial $\eta^2 = 0.64$ | $F(1,6) = 1.08$<br>$p = 0.34$<br>partial $\eta^2 = 0.15$ | $F(11,66) = 0.33$<br>$p = 0.98$<br>partial $\eta^2 = 0.05$ |
| Pooled<br>$n = 21$ | $F_{\epsilon}(4,82) = 12.82$<br>$p < 0.0001^{**}$<br>partial $\eta^2 = 0.39$ | $F(1,20) = 7.89$<br>$p = 0.011^*$<br>partial $\eta^2 = 0.28$ | $F_{\epsilon}(3,56) = 2.49$<br>$p = 0.074$<br>partial $\eta^2 = 0.11$ |

Note—All mice in *Exp.1* (sessions 1–6: FR1; sessions 7–12: FR3) were included in the table, including the mouse that was classified as a non-responder in **Figure 1b**. $^\dagger$ One mouse in *Ymaze.C1* died due to fighting during transportation prior to arrival at the colony (see also Methods: *Animals and housing condition*). Partial $\eta^2$ values of 0.01, 0.06, and 0.14 can be used as a benchmark for small, medium, and large effect sizes, respectively; effect sizes are calculated as $SS_{\text{effect}} / (SS_{\text{effect}} + SS_{\text{error}})$ (Cohen, 1988); yellow boxes: large effect sizes with partial $\eta^2 > 0.14$ ; \*: $p \leq 0.05$ ; \*\*: $p < 0.005$

**Supplementary Table 3:**
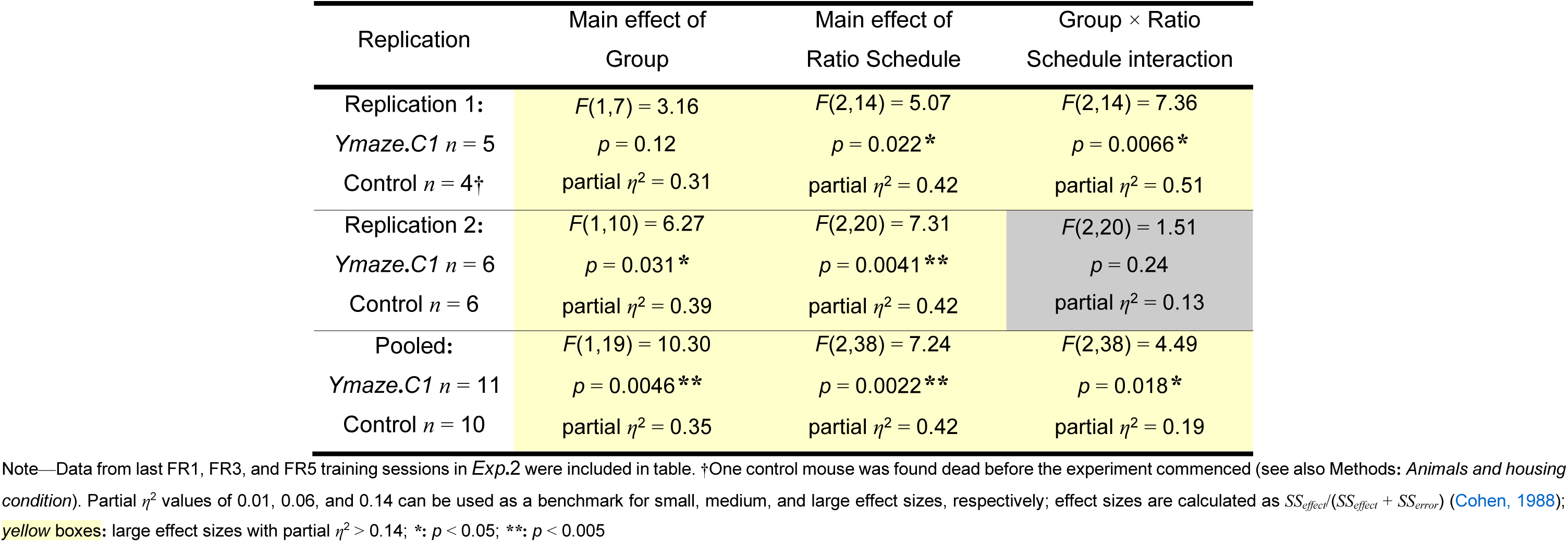

| Replication | Main effect of Group | Main effect of Ratio Schedule | Group × Ratio Schedule interaction |
| --- | --- | --- | --- |
| Replication 1: | $F(1,7) = 3.16$ | $F(2,14) = 5.07$ | $F(2,14) = 7.36$ |
| *Ymaze.C1* $n = 5$ | $p = 0.12$ | $p = 0.022^*$ | $p = 0.0066^*$ |
| Control $n = 4^\dagger$ | partial $\eta^2 = 0.31$ | partial $\eta^2 = 0.42$ | partial $\eta^2 = 0.51$ |
| Replication 2: | $F(1,10) = 6.27$ | $F(2,20) = 7.31$ | $F(2,20) = 1.51$ |
| *Ymaze.C1* $n = 6$ | $p = 0.031^*$ | $p = 0.0041^{**}$ | $p = 0.24$ |
| Control $n = 6$ | partial $\eta^2 = 0.39$ | partial $\eta^2 = 0.42$ | partial $\eta^2 = 0.13$ |
| Pooled: | $F(1,19) = 10.30$ | $F(2,38) = 7.24$ | $F(2,38) = 4.49$ |
| *Ymaze.C1* $n = 11$ | $p = 0.0046^{**}$ | $p = 0.0022^{**}$ | $p = 0.018^*$ |
| Control $n = 10$ | partial $\eta^2 = 0.35$ | partial $\eta^2 = 0.42$ | partial $\eta^2 = 0.19$ |

Note—Data from last FR1, FR3, and FR5 training sessions in *Exp.2* were included in table. $^\dagger$ One control mouse was found dead before the experiment commenced (see also Methods: *Animals and housing condition*). Partial $\eta^2$ values of 0.01, 0.06, and 0.14 can be used as a benchmark for small, medium, and large effect sizes, respectively; effect sizes are calculated as $SS_{effect}/(SS_{effect} + SS_{error})$ (Cohen, 1988); **yellow boxes**: large effect sizes with partial $\eta^2 > 0.14$ ; \*: $p < 0.05$ ; \*\*: $p < 0.005$

**Supplementary Table 4:**
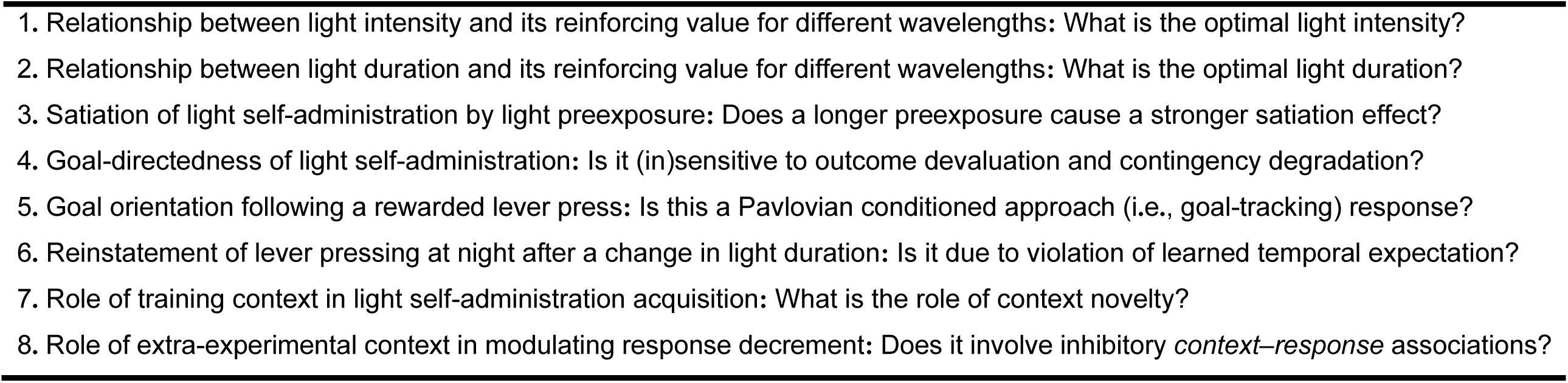

